# Human plastins are novel cytoskeletal pH sensors with a reduced F-actin bundling capacity at basic pH

**DOI:** 10.1101/2025.03.26.645573

**Authors:** Lucas A. Runyan, Elena Kudryashova, Richa Agrawal, Mubarik Mohamed, Dmitri S. Kudryashov

**Author notes:** Correspondence to Dmitri S. Kudryashov. Brandeis University, Waltham, MA, USA, 02453. Versatope Therapeutics, Lowell, MA, USA 01852. University of Utah School of Medicine, Department of Ophthalmology, Salt Lake City, UT, USA, 84132.

## Abstract

Intracellular pH (pH_i_) is a fundamental component of cell homeostasis. Controlled elevations in pH_i_ precede and accompany cell polarization, cytokinesis, and directional migration. pH dysregulation contributes to cancer, neurodegenerative diseases, diabetes, and other metabolic disorders. While cytoskeletal rearrangements are crucial for these processes, only a few cytoskeletal proteins, namely Cdc42, cofilin, talin, cortactin, α-actinin, and AIP1 have been documented as pH sensors. Here, we report that actin-bundling proteins plastin 2 (PLS2, aka LCP1) and plastin 3 (PLS3) respond to physiological scale pH fluctuations by a reduced F-actin bundling at alkaline pH. The inhibition of PLS2 actin-bundling activity at elevated pH stems from the reduced affinity of the N-terminal actin-binding domain (ABD1) to actin. In fibroblast cells, elevated cytosolic pH caused the dissociation of ectopically expressed PLS2 from actin structures, whereas acidic conditions promoted its tighter association with focal adhesions and stress fibers. We identified His207 as one of the pH-sensing residues whose mutation to Lys and Tyr reduces pH sensitivity by enhancing and inhibiting the bundling ability, respectively. Our results suggest that weaker actin bundling by plastin isoforms at alkaline pH favors higher dynamics of the actin cytoskeleton. Therefore, like other cytoskeleton pH sensors, plastins promote disassembly and faster dynamics of cytoskeletal components during cytokinesis and cell migration. Since both plastins are implemented in cancer, their pH sensitivity may contribute to the accelerated proliferation and enhanced invasive and metastatic potentials of cancer cells at alkaline pH_i_.

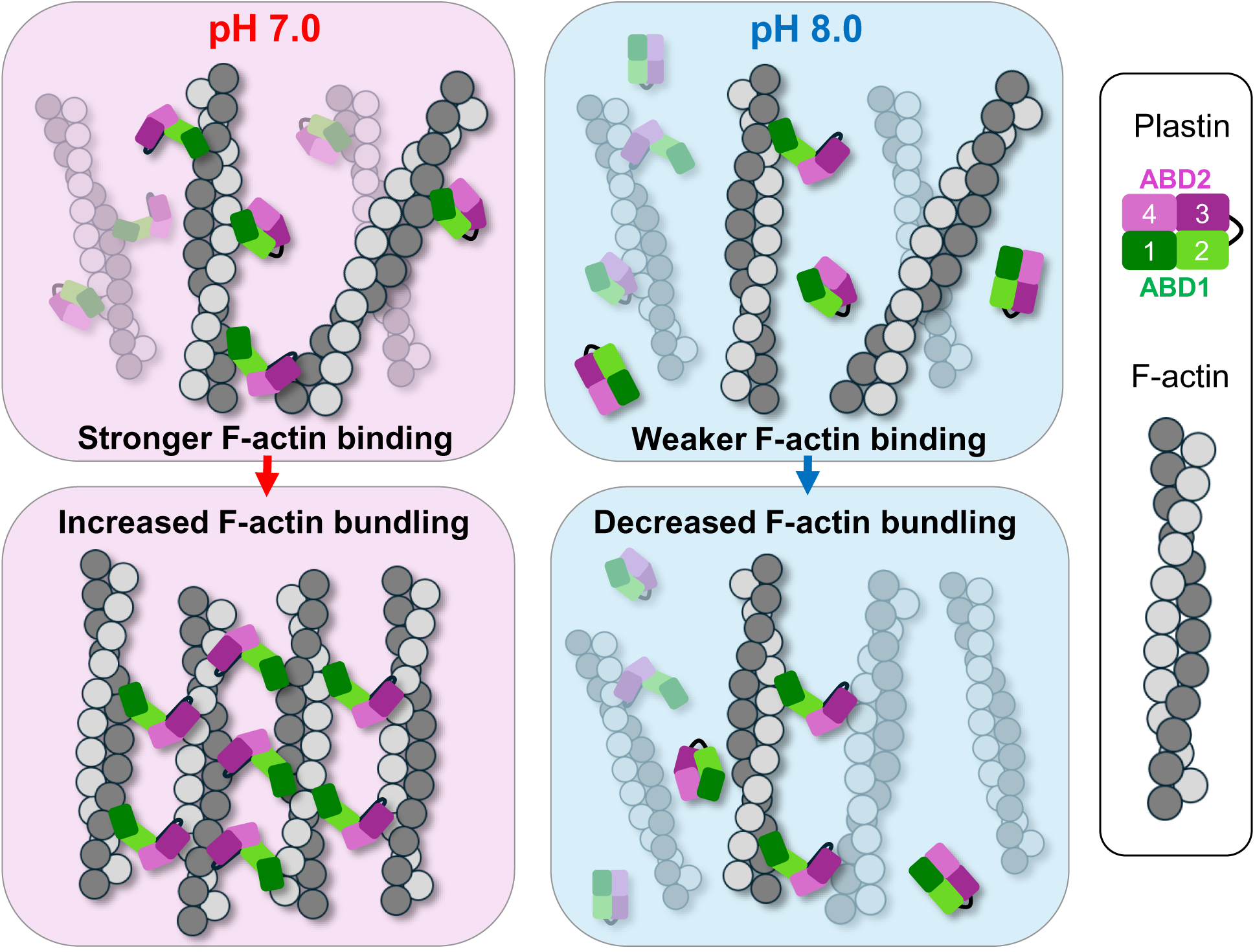

## Introduction

Intracellular pH (pH_i_) is a fundamental parameter of homeostasis. While in most human cells, pH_i_ is maintained at ∼7.2,^1^ the exact pH_i_ value varies for different cells and different stages of the cell cycle, raising up to 7.8-8.0 during the transition from G2 to mitosis.^2^ pH_i_ homeostasis is frequently distorted under pathological conditions such as cancer, where pH_i_ is constitutively increased,^3^ and neurodegenerative diseases, where pH_i_ is decreased.^4^ pH_i_ is predominantly controlled by Na^+^/H^+^ exchanger 1 (NHE1), which utilizes the Na^+^ gradient to extrude H^+^ into the extracellular space.^5^ This process can create short-lived local changes in pH_i_, such as nanodomains with increased pH_i_ (∼7.7) at focal adhesions (FAs) essential for their physiological turnover.^6–8^ At a larger scale, asymmetric localization and activity of the proton pumps across the cell can produce global pH_i_ gradients essential for cell polarization and directional migration.^9–11^ With more basic pH_i_ at the protruding cell front, characterized by high levels of actin dynamics, and more acidic pH_i_ at the retracting cell rear.^12^

The actin cytoskeleton senses and responds to physiological pH fluctuations through regulatory proteins [*e.g.,* Cdc42^13^ and focal adhesion kinase (FAK)^8^], and structural actin-binding proteins [*e.g.,* cofilin,^14,15^ talin,^16^ cortactin^17^, and actin-interacting protein 1 (AIP1)^18^], whose coordinated activity is essential for facilitating actin dynamics, establishing cell polarity, and promoting migration. At the leading edge, a Rho family GTPase Cdc42 is activated by elevated pH_i_ to promote lamellipodial actin assembly.^19,13^ Actin recycling is further facilitated by elevated pH_i_ via a release of the inhibitory association of cortactin with cofilin,^17^ allowing cofilin, whose actin-severing activity is more potent at elevated pH,^20–22^ to effectively sever actin filaments in lamellipodia.^23^

At the adhesion level, elevated pH_i_ promotes the autophosphorylation of FAK, which initiates the recycling of nascent focal adhesions formed at the leading edge, targeting many of them for disassembly.^8^ Recycling of mature FAs is also favored by elevated pH_i_ via weakening talin’s affinity for F-actin^16^ and promoting cofilin’s severing capacity,^20–22^ causing disintegration of FAs and disassembly and recycling of actin filaments. It is conceivable that other FA components are also regulated by elevated pH_i_. Thus, it has been shown that α-actinins from *Dictyostelium discoideum*^24^ and *Hemicentrotus pulcherrimus*^25^ are inhibited by high pH_i_ in the physiological range. However, whether mammalian α-actinin isoforms have a similar pH sensitivity is not known.

Plastin is an actin-bundling protein enriched at the leading edge and in FAs. Like α-actinins, plastins belong to a large family of tandem calponin-homology (*t-*CH) domain actin organizers.^26^ Plastins (Fig. 1A) contain an N-terminal calcium-binding regulatory domain (RD), followed by two actin-binding domains (ABD1 and ABD2), each composed of two calponin-homology (CH) domains.^27^ In the absence of filamentous actin (F-actin), plastins are arranged in a compact horse-shoe-like arrangement^28^. In this form, the ABDs are engaged in an inhibitory association, which permits weak binding to F-actin via one of the domains but does not permit bundling.^27,29,30^ Upon binding to F-actin, plastins undergo a structural rearrangement that separates the ABDs, allowing them to bridge actin filaments, cross-linking them into bundles and networks.^29,30^ These assemblies are components of cellular structures such as filopodia, lamellipodia, FAs, podosomes, and the cortical cytoskeleton.^31,32^ Of the three human plastin isoforms, PLS1 (I-plastin, fimbrin) is expressed in the inner ear stereocilia and intestinal microvilli,^33–36^ PLS2 (L-plastin, LCP1) in immune cells,^33,34,37^ and PLS3 (T-plastin) in all solid tissues.^33,34,38^ Importantly, PLS2 is ectopically expressed in many cancers, and its expression correlates with enhanced invasive and metastatic potential^39–41^, in agreement with its localization to invadopodia and the leading edge.^42^ PLS3 contributes to carcinogenesis by fostering drug resistance,^43^ an activity which has also been suggested for PLS2 in myelomas.^44^ PLS3 has also been implicated in promoting migration, invasiveness, and proliferation of cancer cells.^45,46^

**Figure 1.**
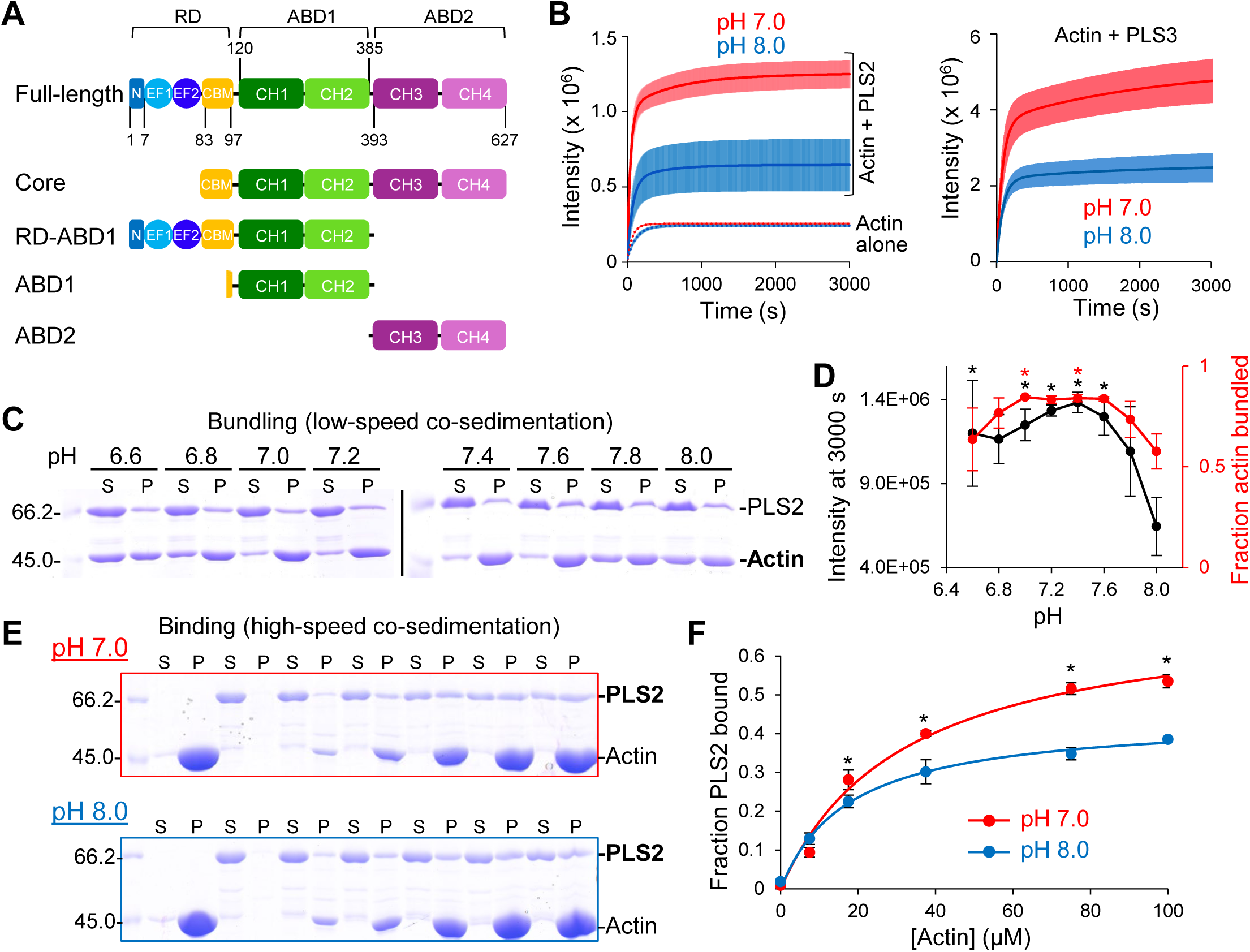
PLS2 bundling ability is negatively regulated by alkaline pH. (A) Schematic of plastin domains and PLS2 constructs used in this study. N, N-terminal 7-a.a. sequence; EF1 and 2, EF-hand motifs; CBM, calmodulin-binding motif; CH1-4, calponin-homology domains; RD, regulatory domain; ABD1 and 2, actin-binding domains. Numbers represent PLS2’s amino acid sequence. (B) Light scattering traces of F-actin bundling by PLS2 (left panel; full range of traces at pH from 6.6 to 8.0 is shown in Fig. S1A) and that of by PLS3 (right panel) at pH 7 (red) and pH 8 (blue). Dotted lines (left panel) represent light scattering traces upon actin polymerization at pH 7.0 (red) and pH 8.0 (blue); see also Fig. S1E. Solid lines represent averaged extrapolated data; colored areas represent SD of the mean (*n* ≥ 3). (C) Representative 10% SDS-PAGE gels of supernatant (S) and pellet (P) fractions from low-speed co-sedimentation of actin bundles formed by PLS2 under different buffer pH conditions. (D) Light scattering intensity (black) of the reactions at 3000 s (shown in B and in Fig. S1A) and the fraction of actin pelleted (red) in low-speed centrifugation assays (shown in C) were plotted as functions of pH. Error bars represent the SD of the mean (n ≥ 3); asterisks are color-coded and indicate significant differences from the corresponding value at pH 8.0. *P* values for light-scattering and centrifugation assays are reported in Tables S1 and S2, respectively. (E) Representative 10% SDS-PAGE gels of supernatant (S) and pellet (P) fractions from high-speed co-sedimentation of PLS2 with increasing concentrations of F-actin at pH 7.0 and 8.0. (F) Quantitation of the high-speed co-sedimentation data (E). Error bars represent SD of the mean (*n* = 3). Asterisks indicate significant difference in fraction of PLS2 bound to actin at 17.5 (*P* = 0.014), 37.5 (*P* = 0.04), 75 (*P* = 0.014), and 100 µM actin (*P* = 0.005).

The actin-bundling activity of plastins is inhibited by Ca^2+^ and enhanced by phosphorylation.^27,29,47^ The inhibition by Ca^2+^ is well known, albeit its mechanism remains obscure as the location of the regulatory, Ca^2+^-binding domain (RD) relative to the actin-binding core is not structurally resolved and only indirectly implied at the ABD1-ABD2 interface.^48^ Upon initial binding of two Ca^2+^ ions, the EF-hands in the RD interact with a linker region containing a calmodulin-binding motif (CBM), reorganizing it into an α-helix.^27,49^ The RD inhibits F-actin binding by one of the ABDs, whose identity is debated.^27,29,30^ The phosphorylation-dependent regulation mechanisms of PLS2 are diverse.^42,47,50–55^ Phosphorylation of Ser5 stimulates F-actin bundling in cells^42,47^ but a phosphomimetic mutation of PLS2, S5D, does not appear to change its behavior *in vitro*,^27^ suggesting the possible participation of unknown cellular partners. Identified as a common phosphorylation site in high-throughput studies,^50–54^ Ser406 is strategically positioned to control the inhibitory association between ABDs. The S406E phosphomimetic mutation weakens binding between ABD1 and ABD2, releasing the autoinhibition and leading to dramatically potentiated bundling both *in cellulo* and *in vitro,* even in the presence of Ca^2+^.^29^

In this study, we demonstrate that F-actin bundling by PLS2 and PLS3 is pH-dependent, with weakened bundling activity at basic pH. We found that PLS2’s pH-sensitivity derives from the inhibited F-actin-binding ability of ABD1 at basic pH, while actin binding by ABD2 was less affected by pH. We validated the physiological importance of pH-sensing by PLS2 through two independent methods. First, by changing the pH_i_ of fibroblasts, we found that PLS2 redistributed from mainly F-actin-associated to diffuse cytosolic upon transition from neutral to basic pH. Second, we generated mutants of PLS2 whose actin-bundling activity was pH-independent. These mutants allowed us to show the cellular consequences of the loss of pH-sensitivity of PLS2, thereby proving that pH-sensitivity is a *bona fide* sensory mode of PLS2, with physiologically relevant consequences to its activity and distribution in cells. This study suggests that the pH sensitivity of PLS2 and PLS3 likely contributes to the reported enhanced migration, invasion, and proliferation of cancer cells expressing these plastin isoforms.

## Results

### Human plastin isoforms PLS2 and PLS3 bundle F-actin in a pH-dependent manner

We evaluated the pH sensitivity of F-actin bundling by PLS2 and PLS3 via light scattering (Fig. 1B) and by PLS2 via differential centrifugation under conditions sufficient to pellet bundled F-actin but not individual filaments (20,000 x g, 20 min; Fig. 1C,D). F-actin was bundled by PLS2 across all conditions tested, from pH 6.6 to pH 8.0, with the most intense bundling occurring between pH 7.0 and 7.6 as judged from both higher intensity of light scattering and more actin in the pellet at low-speed centrifugation (Fig. 1B-D; S1A; Tables S1, S2). At pH 7.8 and 8.0, F-actin bundling by PLS2 was progressively decreased (Fig. 1D). Similarly, PLS3 bundled F-actin at pH 7.0 more efficiently than at pH 8.0 (Fig. 1B, right panel). These data suggest that PLS2 and PLS3 bundle F-actin in a pH-dependent manner. Since both plastin isoforms showed similar pH dependence, we focused on the characterization of PLS2 in the subsequent experiments to streamline the experimental process.

Next, PLS2’s binding to F-actin was measured at pH 7.0 and 8.0 via high-speed co-sedimentation (300,000 x g, 30 min), where the centrifugation speed is sufficient to co-pellet all F-actin together with bound proteins. PLS2 co-sedimented with F-actin significantly more effectively at pH 7.0 than at pH 8.0 (Fig. 1E,F), in good agreement with light scattering and low-speed co-sedimentation bundling data (Fig. 1B-D).

F-actin bundling by plastin is sensitive to ionic strength.^56^ Since the same 10 mM HEPES buffer was used to create buffers with different pHs, the increased concentration of NaOH added to the pH 8.0 buffer contributed additional ionic strength equivalent to an extra 6 mM KCl (Supplementary Methods).^57^ To ensure that the inhibited actin bundling behavior of PLS2 at pH 8.0 is not an artifact of the increased ionic strength, F-actin bundling by PLS2 was re-assessed at pH 7.0 with 36 mM KCl (up from 30 mM KCl) and at pH 8.0 with 24 mM KCl (down from 30 mM KCl). We found that the 6 mM difference in KCl concentration had a negligible effect on actin bundling by PLS2, as reported by light scattering intensity and actin pelleting in low-speed co-sedimentation assays (Fig. S1B-D).

Another potential artifact could arise from the pH sensitivity of actin polymerization *per se*, with acidic pH promoting polymerization and basic pH inhibiting it.^58,59^ To rule out the possibility that low light scattering intensity reflects the differences in polymerization rather than bundling, we compared the light scattering intensity of F-actin polymerization at pH 7.0 and 8.0. While the actin polymerization rate was slightly slower at pH 8.0 compared to pH 7.0 (Fig. 1B, actin alone; S1E), the final light scattering intensity for each pH condition was nearly identical (Fig. S1E), and the extent of F-actin pelleted at high-speed centrifugation was the same for both pH conditions (Fig. S1F,G). These observations agree with the reported minor differences in the critical concentration of F-actin between pH 7.0 and 8.0.^58^ Therefore, the pH dependence of actin polymerization is not the source of decreased actin-bundling activity of PLS2 at elevated pH.

The 7.8-8.0 pH range, required for inhibition of F-actin bundling by PLS2, is at the upper border of the physiological range, as these values are reached during the G2/M transition of the cell cycle^2^ and the local pH_i_ near FAs^8^ supporting the role of PLS2 as a physiologically relevant pH-dependent actin regulator.

### The PLS2 actin-binding core retains pH sensitivity

The role of individual plastin domains in the observed pH sensitivity of actin bundling was evaluated using PLS2 truncation constructs (Fig. 1A). A possible involvement of the regulatory calcium-binding domain was tested using PLS2-core (a.a. 83-627), a construct containing both ABDs, but missing the Ca^2+^-binding EF-hands. We chose to include residues N-terminal to CH1 in the construct, as the analogous residues are implicated in actin binding in other members of the *t-*CH protein superfamily.^60,61^ PLS2-core bundled F-actin weaker than the full-length (FL) protein, with lower maximum light scattering intensity and less actin pelleted under both pH conditions tested (Figs. 1B-D; 2A-C). Yet, basic pH inhibited PLS2-core’s F-actin bundling ability by a factor of ∼2 (Fig. 2A-C), suggesting that the PLS2-core construct missing the RD is still sensitive to pH, similar to the FL PLS2.

**Figure 2.**
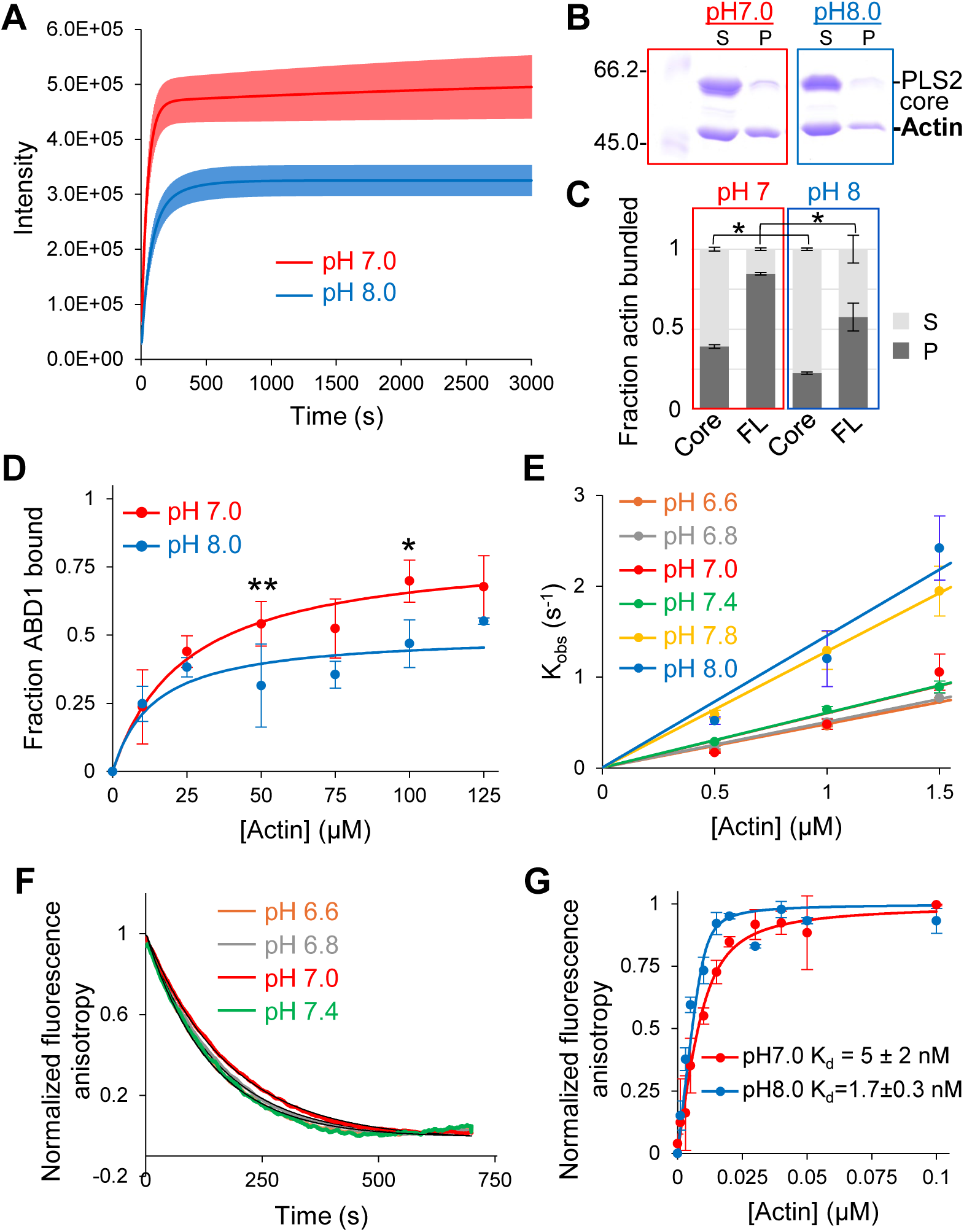
pH-dependence of PLS2 stems from weakened ABD1 binding to actin at basic pH. (A) Light scattering traces of F-actin bundling by PLS2-core at pH 7.0 (red) and 8.0 (blue). Solid lines represent averaged extrapolated data; colored areas represent SD of the mean (*n =* 3). (B) Representative 10% SDS-PAGE gels of supernatant (S) and pellet (P) fractions from low-speed centrifugation of actin bundles formed by PLS2-core at pH 7.0 and 8.0. (C) Quantitation of the low-speed centrifugation data; error bars represent SD of the mean (*n* = 3; *, *P* = 0.01). (D) Fraction of ABD1 depleted from the supernatant during high-speed co-sedimentation with increasing F-actin concentrations at pH 7.0 (red) and pH 8.0 (blue). Error bars represent SD of the mean (*n* = 3, except for 50 and 100 µM Actin points, where *n* = 6; **, *P* = 0.002; *, *P* a= 0.02). (E) Slopes from linear fits of *K_obs_* values (symbols) determined from kinetic association experiments of FM-ABD2 with F-actin were used to obtain *k_on_* values for each pH value (Table 1); error bars represent SD of the mean (*n* ≥ 8). (F) Dissociation kinetics (color curves) of FM-ABD2 from F-actin upon competition with excess unlabeled ABD2 were averaged (n≥4) and fit to single exponential (black curves) to obtain *k_off_* values (Table 1). (G) Equilibrium binding data (symbols) of FM-ABD2 binding to phalloidin-stabilized F-actin at pH 7 (red) and pH 8 (blue) fit to a quadratic isotherm (lines). Error bars represent the SD of the mean (*n* = 3). K_d_ values are the mean ± SD of three technical replicates.

**Table 1.**
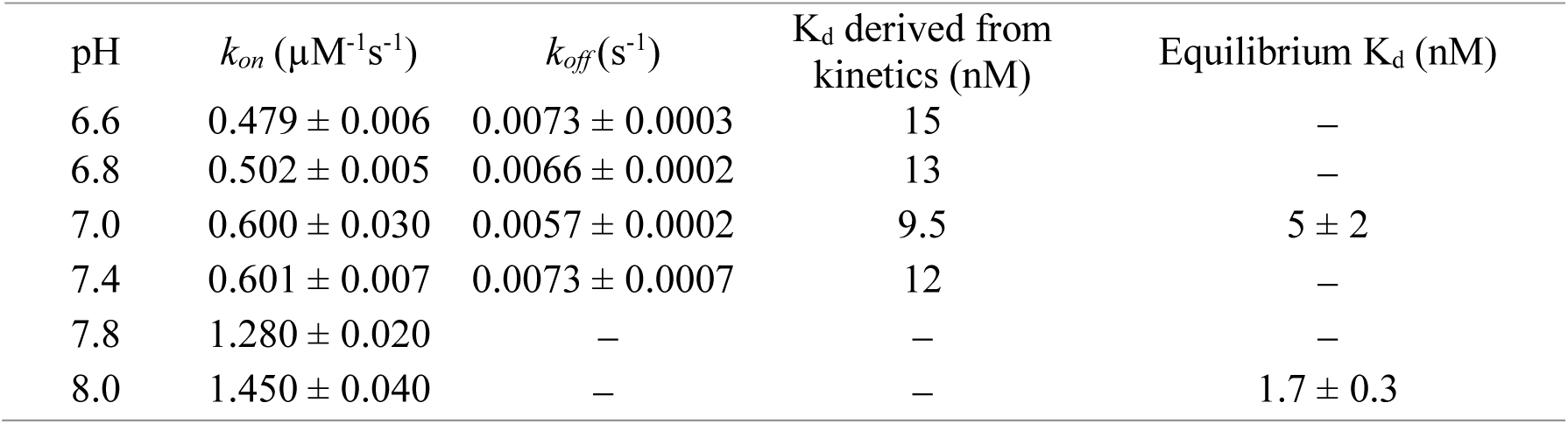
Kinetic and equilibrium parameters for association and dissociation of FM-ABD2 to and from F-actin at varying buffer pH. *k_on_* (n ≥ 8) and *k_off_* (n ≥ 4) and equilibrium K_d_ (n = 3) values are reported as mean ± SD. K_d_ derived from kinetics was calculated using the *k_off_* and *k_on_* for each specified pH.

| pH | $k_{on}$ ($\mu\text{M}^{-1}\text{s}^{-1}$) | $k_{off}$ ($\text{s}^{-1}$) | $K_d$ derived from kinetics (nM) | Equilibrium $K_d$ (nM) |
| --- | --- | --- | --- | --- |
| 6.6 | $0.479 \pm 0.006$ | $0.0073 \pm 0.0003$ | 15 | — |
| 6.8 | $0.502 \pm 0.005$ | $0.0066 \pm 0.0002$ | 13 | — |
| 7.0 | $0.600 \pm 0.030$ | $0.0057 \pm 0.0002$ | 9.5 | $5 \pm 2$ |
| 7.4 | $0.601 \pm 0.007$ | $0.0073 \pm 0.0007$ | 12 | — |
| 7.8 | $1.280 \pm 0.020$ | — | — | — |
| 8.0 | $1.450 \pm 0.040$ | — | — | $1.7 \pm 0.3$ |

### pH-dependence of PLS2 stems from weakened ABD1 binding to actin at basic pH

To test whether actin-binding abilities of individual ABDs of plastins are pH-dependent, we focused on ABDs of PLS2, as ABD2 of PLS3 is unstable and cannot be isolated.^27^ While RD contributes to the association between ABD1 and ABD2, its contribution to the binding of ABD1 to actin is insubstantial.^27^ Furthermore, RD-ABD1 (a.a. 1-385) has similar molecular weight to actin, which complicates the SDS-PAGE analysis. Therefore, we used ABD1 (a.a. 120-385), which binds F-actin with an affinity similar to that of RD-ABD1 but migrates on SDS-PAGE substantially faster.^27^ In high-speed co-sedimentation assays, consistently less ABD1 co-pelleted with F-actin at pH 8.0, as compared to pH 7.0 at all tested actin concentrations (Fig. 2D). Since the statistical significance in triplicates was borderline, we conducted another round of triplicates at actin concentrations 50 and 100 µM. Data combined from the two experiments for these two actin concentrations indeed reached the significance level (Fig. 2D). Note that this inhibition at basic pH is consistent with the inhibited interactions of FL PLS2 and PLS2-core with F-actin.

We noticed that only a fraction of ABD1 was found in the pellet (*i.e.*, bound to F-actin) after high-speed co-sedimentation, despite the large excess of actin and apparent saturation of the binding curves at both pH conditions (Fig. 2D). This behavior was similar to that of FL PLS2 (Fig. 1E,F), except that the latter, but not the former, could also result from actin bundling by the FL protein, which is more difficult to interpret. We hypothesized that the apparent saturation at low binding levels of ABD1 may reflect the limitation of the co-pelleting approach, which does not truly report the binding equilibrium. Indeed, the duration of pelleting required to separate the F-actin complexes from free proteins upon centrifugation is sufficient for the dissociation of weakly bound complexes. Therefore, the method may not adequately report the dissociation constants (K_d_), particularly for reactions characterized by fast dissociation rate constants (*k_off_*). We confirmed that the incomplete binding is not due to the protein’s structural or functional inadequacy by re-pelleting the supernatant with more added F-actin and demonstrating a similar distribution of ABD1 between pellet (bound to F-actin) and supernatant (unbound) fractions as in the original experiment (Fig. S1H).

We next evaluated the pH sensitivity of ABD2 binding to actin. Since ABD2 binds to F-actin with nanomolar K_d_,^29^ we took advantage of the sensitivity of fluorescence anisotropy to analyze the pre-steady state binding and dissociation kinetics of fluorescein-maleimide labeled ABD2 (FM-ABD2) using a stopped-flow fluorometer. Interestingly, while the *k_on_* values of FM-ABD2 binding to F-actin changed little in the pH 6.6 −7.4 range, they increased 2- to 3-fold upon transition to pH 7.8 and 8.0 (Fig. 2E; Table 1). The *k_off_* values also remained nearly constant in the 6.6 to 7.4 range of pH (Fig. 2F), resulting in K_d_ values fluctuating unsubstantially in the 10-15 nM range (Table 1). We could not measure *k_off_* at pH 7.8 and 8.0 due to the low signal-to-noise ratio for reasons we do not fully understand, rendering the data uninterpretable. To address this shortage, the affinity of FM-ABD2 to F-actin was measured independently in equilibrium anisotropy experiments at pH 7.0 and 8.0 (Fig. 2G) with phalloidin-stabilized actin to prevent its depolymerization at concentrations below critical. The K_d_ of FM-ABD2 for phalloidin-stabilized F-actin at pH 7.0 was 5 ± 2 nM, *i.e.,* within a reasonable margin of 9.5 nM measured in the kinetic assay.^62^ At pH 8.0, the K_d_ of FM-ABD2 binding to F-actin was 1.7 ± 0.3 nM, *i.e.,* ∼3-fold stronger than at pH 7.0. Therefore, this difference is mainly accounted for by the higher *k_on_* at basic pH, implying that the *k_off_* of ABD2 from actin is not substantially affected in the pH 6.6-8.0 range. To summarize, ABD2 binds F-actin stronger at basic pH, in contrast to the behavior of both the FL protein and ABD1, supporting our previous observations that the initial interaction of plastin with actin, which promotes conformational changes required for bundling, is mediated via ABD1.^29^

### Inhibitory ABD1-ABD2 association is not responsible for the weaker bundling ability of PLS2 at basic pH

The inhibitory association between ABDs is a key regulatory mechanism in plastins.^29,30^ To test whether the detected pH sensitivity is related to this mutual inhibition, we measured the effect of pH on the binding kinetics of RD-ABD1 and ABD2 (Fig. 3A-C). The dissociation rates of the complex, *k_off_*, did not change significantly across the explored pH conditions, fluctuating around 0.008 s^-1^ (Fig. 3B, Table S3). The association rates slowed moderately at higher pH, reaching ∼2-fold difference between pH 6.6 and 8.0 (Fig. 3A,B, Table S3). K_d_ derived from kinetic rates reasonably agreed with those measured directly in equilibrium experiments, revealing ∼2-fold stronger association at pH 7.0 than pH 8.0 (Fig. 3C, Table S3). While the interaction between RD-ABD1 and ABD2 is mildly pH-sensitive, it favors weakening the inhibitory ABD1-ABD2 association^29^ at the elevated pH (Table S3). If this were the mechanism primarily responsible for the pH sensitivity of PLS2, it should result in stronger bundling at basic pH, which is not the case. Therefore, the observed pH sensitivity of the ABD1-ABD2 association is unlikely to account for the weaker bundling capacity of PLS2 at basic pH.

**Figure 3.**
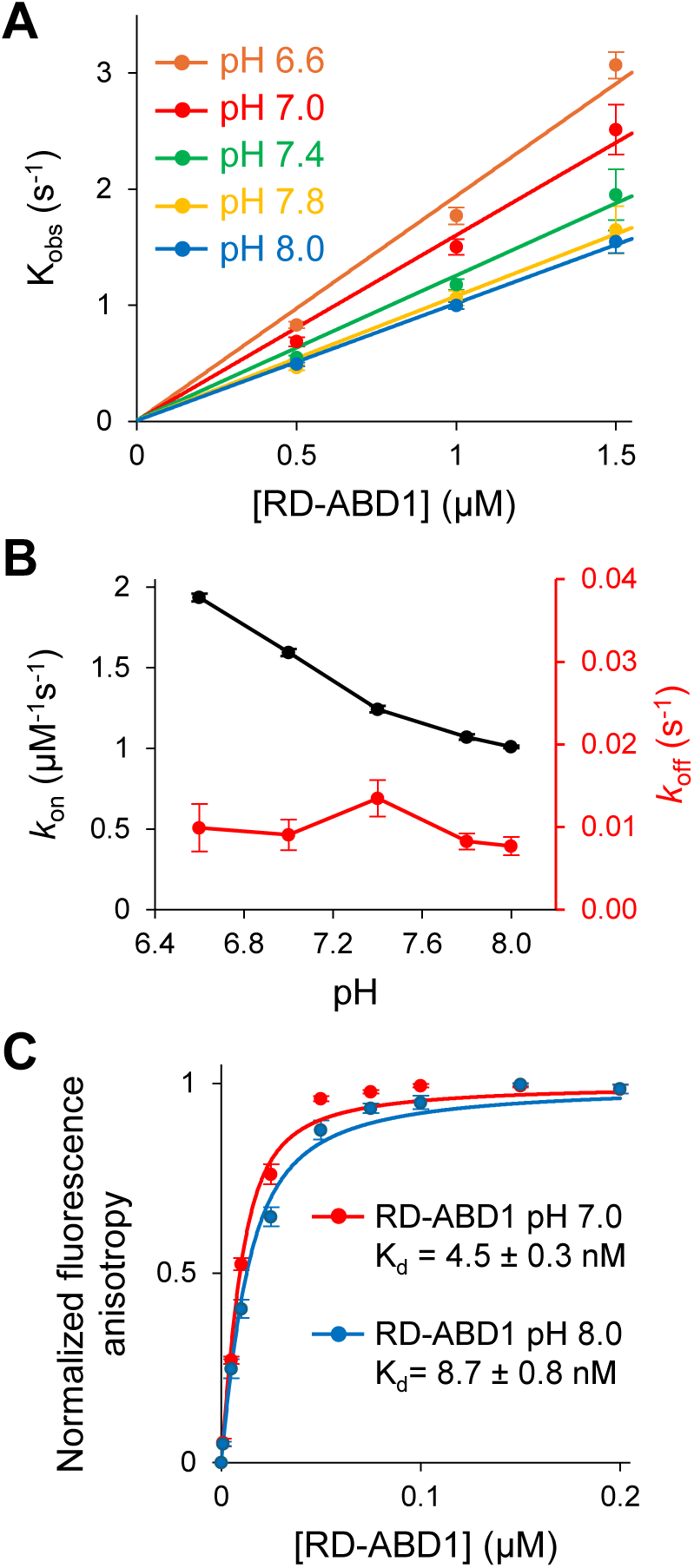
Inhibitory ABD1-ABD2 association is not responsible for the weaker bundling ability of PLS2 at basic pH. (A) Slopes from linear fits of *K_obs_* values (symbols) determined from kinetic association experiments of FM-ABD2 with RD-ABD1 were used to obtain *k_on_* values for various pH conditions (Table S3); error bars represent SD of the mean (*n* ≥ 8). (B) *k_on_* (black) and *k_off_* (red) plotted as a function of pH. *k_on_* values derived from the slope of the linear function fit to experimentally measured *K_obs_* data shown in (A), error bars represent curve fit error. *k_off_* values derived from dissociation experiments of FM-ABD2 from RD-ABD1 with excess unlabeled ABD2, fit to a single exponential. Error bars represent SD of the mean (*n* ≥ 4). (C) Equilibrium binding data (symbols) of FM-ABD2 binding to RD-ABD1 at pH 7.0 (red) and 8.0 (blue) fit to a quadratic isotherm (lines). Error bars represent the SD of the mean (*n* = 3). K_d_ values are reported as the mean ± SD from three technical replicates.

### Screening mutagenesis revealed His207 as a potential pH sensor

Changes in protein properties near neutral pH are commonly mediated by changes in the protonation state of His residues^63^, whose pK_a_ values fall in this range. Histidines are responsible for pH sensitivity of talin, cofilin, FAK, and AIP1.^8,15,16,18^ To explore the role of specific His residues in the pH dependence of F-actin bundling by PLS2, we used His-to-Lys and His-to-Tyr mutations. The former substitution mimics the charge of protonated His (favored by low pH conditions), while the latter mimics the lack of charges of deprotonated His side chain (favored by higher pH), with the reservation that the selected mutations do not well reproduce histidine geometry and other properties.

We created a library of single His-to-Lys replacements for each of the eight His residues in PLS2 and tested their abilities to bundle actin at pH 7 and 8 (Fig. S2). In a preliminary screen with two different ratios of PLS2 to actin, we found that H116K had severely inhibited F-actin bundling, regardless of pH (Fig. S2A). H116 is located in the linker between RD and ABD1 and neighbors Ser117, whose mutation to Glu, S117E, similarly ablates F-actin bundling by PLS2 (Fig. S3). The mechanism by which introducing a charge to His116 or Ser117 inhibits actin bundling is unclear, but the role of this region in actin binding correlates to that of other *t-*CH protein superfamily proteins.^60,61^ While PLS2 constructs carrying H207K, H277K, H378K, and H415K mutations also showed aberrations in the pH sensitivity in the preliminary experiments (Fig. S2A), upon careful characterization at five different ratios of PLS2 to actin, only H207K satisfied the sought pH-insensitivity [*i.e.*, retained bundling ability at basic pH (Fig. S2B)].

### PLS2 H207Y, but not H207K, partially reproduces the mechanisms governing PLS2’s pH-sensitivity

Light scattering and low-speed co-sedimentation experiments showed that PLS2 H207K bundled F-actin effectively and similarly at both pH 7.0 and 8.0, except that the maximum light scattering intensity for H207K at pH 8.0 was even higher than at pH 7.0 (Fig 4A-C). Conversely, H207Y bundled F-actin poorly at pH 7.0 and 8.0 (Fig. 4A-C). Notably, H207Y remained sensitive to pH, as judged by more actin in the pellet and a higher light scattering signal at pH 7.0 than 8.0 (Fig. 4A-C), indicating that residues other than H207 are involved in the pH sensitivity of PLS2.

**Figure 4.**
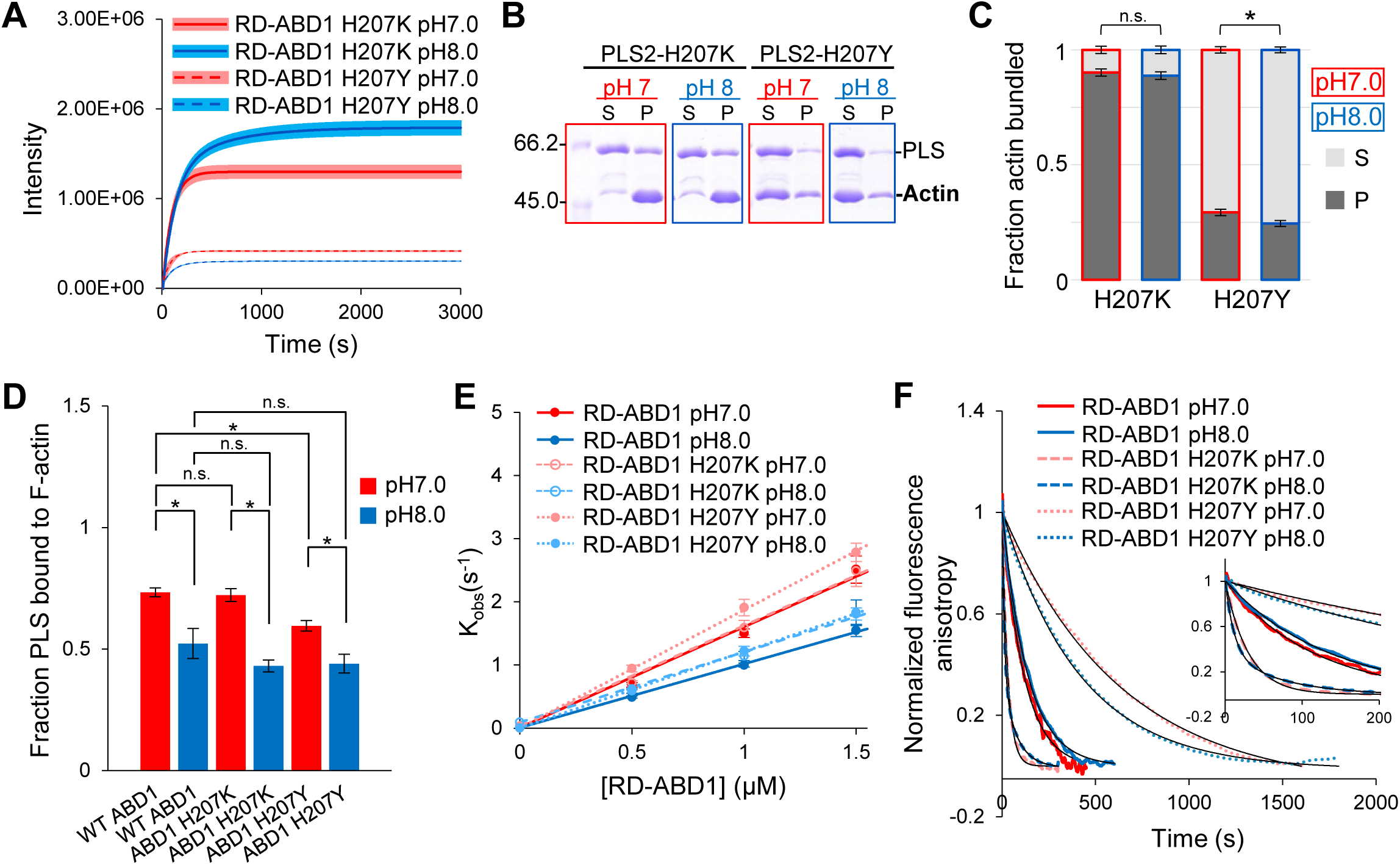
PLS2 H207Y, but not H207K, partially reproduces the mechanisms governing PLS2’s pH-sensitivity. (A) Light scattering traces for F-actin bundling by PLS2 mutants at pH 7.0 (red) and pH 8.0 (blue). Lines are averaged extrapolated data, colored areas are SD of the mean (*n* = 3). (B) Representative 10% SDS-PAGE gels of supernatant (S) and pellet (P) fractions from low-speed co-sedimentation of actin bundles formed by PLS2 mutants at pH 7.0 and 8.0. (C) Quantitation of the low-speed co-sedimentation data. Error bars represent SD of the mean (*n* = 3; *, *P* = 0.003). (D) Fraction of ABD1 constructs pelleted during high-speed co-sedimentation with 100 µM F-actin at pH 7.0 (red) and pH 8.0 (blue). Error bars are SD of the mean (*n* = 3). Significance determined by ANOVA analysis is shown in Table S4. (E) Slopes from linear fits of *K_obs_* values (symbols) determined from kinetic association experiments of FM-ABD2 with the indicated RD-ABD1 constructs were used to obtain *k_on_* values for pH 7.0 and 8.0 (Table S5); error bars represent SD of the mean (*n* ≥ 8). (F) Dissociation kinetics traces (color curves) of FM-ABD2 from the indicated RD-ABD1 were averaged (*n* ≥ 8) and fit to a single or double exponential (black curves) to yield *k_off_* values for each pH condition (Table S5). Inset shows first 200 s of the time-courses for each experiment.

Testing individual ABD1 constructs carrying the same mutations revealed that ABD1-H207Y bound actin weaker than WT ABD1 at pH 7.0 (Fig. 4D), supporting that H207 contributes to tuning the interaction strength in a pH-dependent manner. Yet, at basic pH, both H207Y and H207K constructs bound to F-actin similarly weaker, *i.e.,* demonstrated pH sensitivity similar to that of WT ABD1 (Fig. 4D; Table S4). This observation also suggests that residues other than H207 contribute to this sensitivity. Accordingly, the inability of the H207K mutation to enhance ABD1 binding at basic pH suggests that it potentiates bundling via a different mechanism.

To check whether the enhanced F-actin bundling by H207K may be due to the altered interaction between the actin-binding domains, the *k_on_* and *k_off_* rates of FM-ABD2 interaction with RD-ABD1-H207K and RD-ABD1-H207Y constructs were measured. While RD contributes little to binding of ABD1 to actin,^27^ its contribution to binding to ABD2 is measurable,^29^ justifying its addition to the ABD1 constructs. The mutations had little to no (less than 1.2-fold) effect on the *k_on_* values at both pH 7.0 and 8.0 (Fig. 4E; Table S5), but notably affected the dissociation rate constants, *k_off_* (Fig. 4F; Table S5). Specifically, H207Y decreased *k_off_* by a factor of 3-4, while H207K accelerated the dissociation by a factor of 4-12 (Fig. 4F; Tables S3, S5). The large 4-12 fold gap reflects the double-exponential character of the dissociation rate of RD-ABD1-H207K at pH 8.0 with 62% and 38% amplitudes, suggesting that basic pH enables two distinct binding modes between RD-ABD1-H207K and ABD2 (Fig. 4F).

Taken together, stronger bundling of F-actin by PLS2-H207K at basic pHs stems mainly from the greater rate of dissociation of ABD2 from RD-ABD1. This allows PLS2 to populate the bundling-competent state with weakly connected ABD1 and ABD2 more frequently, resulting in an actin-bundling protein with lower sensitivity to pH. The weaker bundling capacity of PLS2-H207Y stems from two separate mechanisms. First, this mutation inhibits the dissociation of RD-ABD1 from ABD2, strengthening the complex and preventing PLS2 from adopting the bundling-competent state. Second, H207Y lowers the affinity of ABD1 to actin, partially reproducing the pH sensitivity mechanism of WT PLS2. Combined, these effects cause the inhibition of actin bundling by PLS2-H207Y, even at pH 7.0.

### PLS2 association with cellular F-actin structures is enhanced by acidic and neutral pH_i_ and reduced by basic pH_i_

To assess whether PLS2 shows pH-dependent behavior in the cellular context, we performed live-cell imaging of XTC fibroblasts transiently transfected with mEmerald-tagged PLS2 constructs, while manipulating pH_i_ via nigericin clamping^64^ (Fig. 5). The pH_i_ of the cells was evaluated using ratiometric imaging of the genetically encoded pH sensor mCherry-SEpHluorin in nigericin-containing buffers adjusted to specific pH values.^65^ The generated calibration curve was linear in the tested range of pH 6.5-8.0 and confirmed that pH_i_ before clamping (*i.e.,* under normal cell culture conditions in the absence of nigericin) was ∼7.4 (Fig. S4A,B).

**Figure 5.**
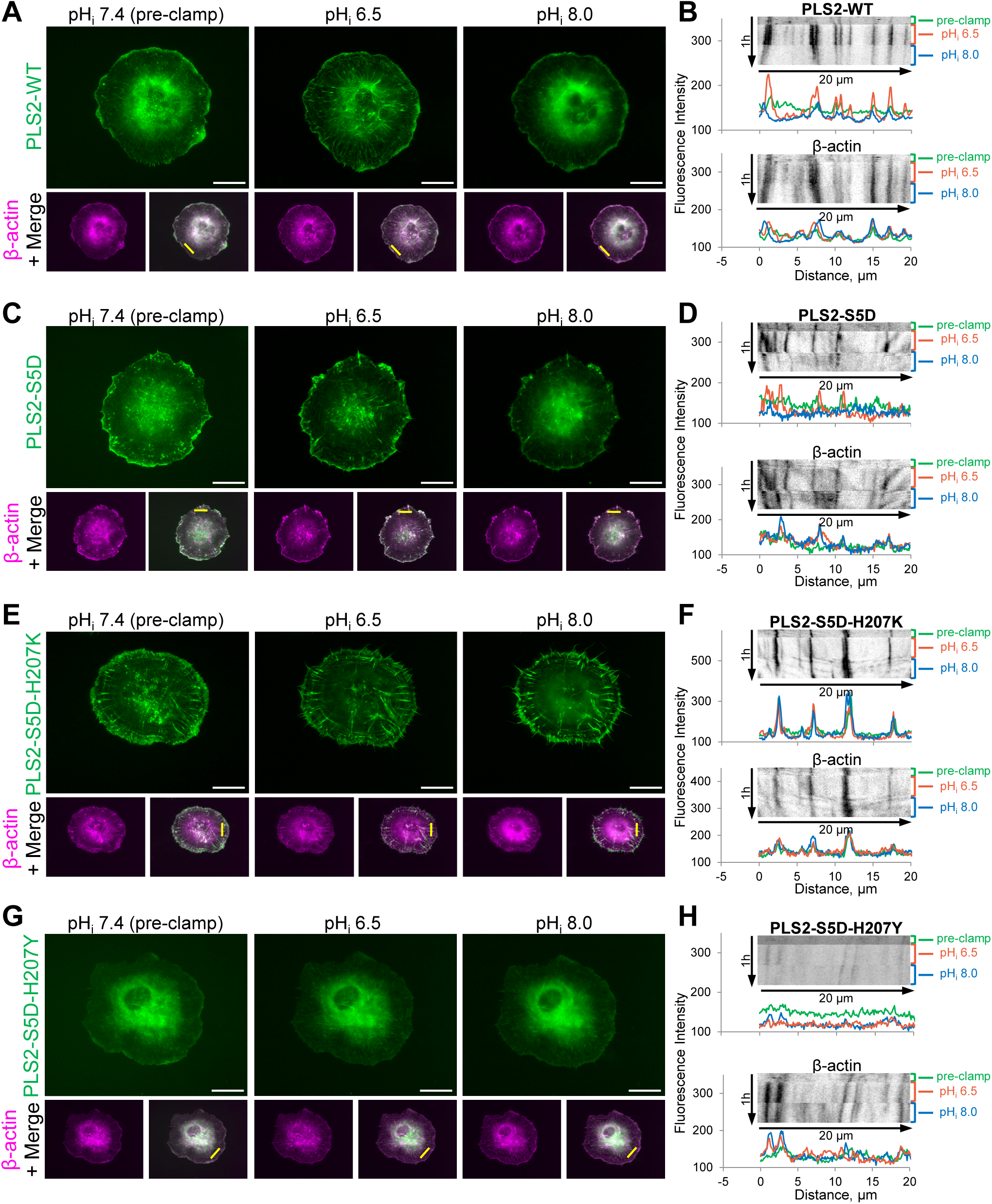
PLS2 shows pH-dependent cellular localization. (A, C, E, F) Time-lapse images of XTC cells co-transfected with mCherry-β-actin (magenta) and mEmerald-tagged WT (A), S5D (C), S5D/H207K (E), or S5D/H207Y (F) PLS2 constructs (green). Cells were imaged pre-clamped and sequentially clamped with nigericin buffers at pH 6.5 and 8.0. First frames of the time-lapse series at each pH_i_ condition (pre-clamping pH_i_ 7.4; pH_i_ 6.5; pH_i_ 8.0) are shown. Scale bars are 20 µm. (B, D, F, H) Kymographs and line plot profiles (taken from the regions indicated in the merge images in A, C, E, F by yellow lines, which also serve as 20 µm scale bars) are shown in two channels (for the corresponding PLS2 construct and β-actin). Plot profiles (*i.e.*, fluorescence intensity traces plotted versus distance indicated by the yellow lines in A, C, E, F at each pH_i_ condition) are color-coded for each pH_i_ according to the color scheme on the kymographs. See also Videos S1-S4.

Under pre-clamping conditions, both WT and phosphomimic S5D PLS2 showed diffuse cytosolic localization with enrichment at the cell edge, while also weakly localized at FAs and stress fibers (Fig. 5A-D, S4C; Videos S1,2) in agreement with our previous findings.^48^ When pH_i_ was decreased to 6.5 via nigericin clamping, the majority of PLS2 was redistributed to FAs and stress fibers. Conversely, when pH_i_ was increased to 8.0, PLS2’s association with cellular F-actin structures decreased. Importantly, the observed effects of PLS2 redistribution are not due to the global actin cytoskeleton rearrangements, as mCherry-β-actin distribution and morphology were only marginally affected by the changes in pH (Fig. 5A-D; Videos S1,2).

We also tested whether weakening the pH sensitivity via H207K or H207Y mutations affects the cellular localization of PLS2. H207K and H207Y PLS2 variants reproduced the cellular localization of WT and S5D PLS2 at low and high pH_i_, respectively, while remaining largely insensitive to pH (Fig. 5E-H, S4D; Videos S3,4). In agreement with its higher actin-bundling capacity, H207K strongly decorated F-actin cellular structures under each tested pH_i_ condition (Fig. 5E,F, S4D; Video S3). H207Y showed mostly diffuse distribution with negligible pH-independent co-localization with actin-rich structures (Fig. 5G,H; Video S4). These data exclude a potential pH-sensitive contribution of the mEmerald tag and confirm that the pH-dependent redistribution of WT and phosphomimetic S5D PLS2 in nigericin-clamped cells under acidic and basic pH can indeed be attributed to the pH-sensing properties of PLS2.

To summarize, under cellular conditions, PLS2 mainly recapitulates its pH-regulated behavior observed *in vitro*: neutral pH_i_ increases the association of PLS2 with F-actin, while alkalization of pH_i_ decreases it (Fig. 6A). Minor differences in the pH-dependency at acidic pH under the *in vivo* and *in vitro* conditions, could be attributed to tuning the PLS2 activity by cellular partners and/or post-translational modifications. Together, our data show that PLS2 and PLS3 are *bona fide* pH-sensing proteins with tunable actin-bundling activity in the physiologically relevant pH range.

**Figure 6.**
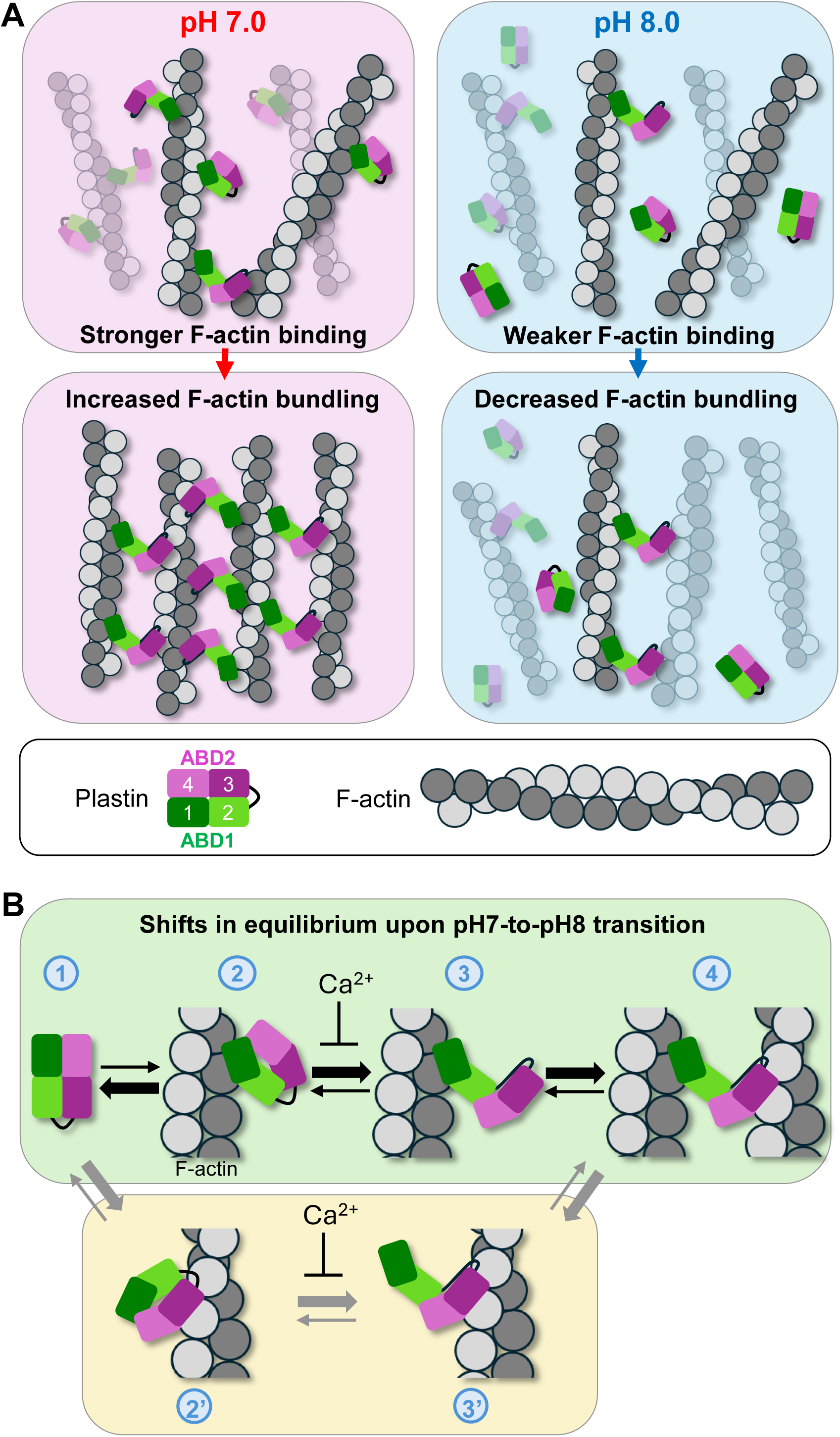
Hypothetical model of pH effects on F-actin bundling by plastins. (A) Plastin’s ability to bundle F-actin is regulated by pH: by respectively promoting and inhibiting the binding of ABD1 to actin (top panels), neutral pH enhances F-actin bundling by PLS, while basic pH reduces bundling (bottom panels). (B) Shifts (represented by thick arrows) in equilibrium states of plastin upon interaction with F-actin during the transition from pH 7.0 to pH 8.0. Transition from free (state 1) to actin-bound state via ABD1 (state2) is disfavored by basic pH. This binding favors the opening of the ABD core (favored by basic pH), enabling ABD2 to sample the space and bind another actin filament (state 3; favored by basic pH), resulting in bundling (state 4). In the alternative scenario (route 1-2’-3’-4), ABD2 binds first (state 2’; favored by basic pH), leading to a partial release of the ABD1-ABD2 inhibition (state 3’, favored by basic pH), followed by ABD1 binding to actin, forming bundles (state 4; disfavored by basic pH). Notice, however, that this latter scenario implies a stronger binding of PLS to actin at alkaline pH, which is not supported by our experimental data (Fig. 1F).

## Discussion

In this study, we introduce PLS2 and PLS3 as novel pH-sensitive cytoskeletal proteins with strongly inhibited F-actin bundling activity at basic pH (Fig. 1B, 6A). We observed a sharp transition towards inhibition of actin bundling by PLS2 at pH > 7.6 (Fig. 1B-D). These pH values are substantially higher than the typical cytosolic values of ∼7.2-7.4 but similar to the higher local pH values measured at focal adhesions (∼7.5 – 7.7)^6,8^ and global pH_i_ measured during the G2/M transition of the cell cycle (∼7.6 – 7.9).^2^ In both processes, the actin cytoskeleton undergoes large-scale rearrangements that likely involve many actin-binding proteins. The increased cytosolic pH coinciding with these rearrangements is consistent with the known roles of talin, cofilin, FAK, cortactin, and AIP1 in favoring actin disassembly and promoting actin dynamics at basic pH.^8,15–18,24^ Therefore, the inhibition of actin bundling by PLS2 and PLS3 at basic pH is consistent with plastins participating in changes in actin dynamics during cell migration and cell division.

The current understanding of the mechanism of F-actin bundling by plastins is summarized in Figure 6B. In solution, ABDs of plastins exist in a tight autoinhibited association (Fig. 6B, state 1) characterized by a nanomolar affinity even when the domains are mixed *in trans*^29^. Despite that ABD2 affinity to actin is also very high (in a low nanomolar K_d_ range),^29^ due to the above ABDs’ autoinhibition, FL plastins interact with actin weakly, and the interaction is primed via only one of the domains, whose identity is debated (states 2 and 2’).^27,29,30,49^ Binding of Ca^2+^ to EF-hands affects the RD interaction with the loop region between ABD1 and ABD2,^48^ either preventing the release of the autoinhibition or sterically blocking the association of the secondary ABD with actin [transition between states 2 (2’) and 3 (3’)]. In the absence of Ca^2+^, the interaction between actin and the primary ABD weakens the autoinhibition,^29^ moderately promoting domain separation, and thus enabling the other ABD to sample the space in search for another actin filament [states 3 (3’)], resulting in bundling (state 4). Our kinetic and equilibrium characterization of PLS2 revealed that basic pH *1)* weakens the interaction of ABD2 with RD-ABD1 by a factor of ∼2 (Fig. 3; Table S3), *2)* enhances the interaction of ABD2 with F-actin by ∼3-fold (Fig. 2E-G; Table 1), and *3)* reduces the interaction of ABD1 with F-actin (Fig. 2D). Since the first two effects should potentiate bundling (*i.e.,* by favoring the transition through states 2’ and 3’ to 4), the observed reduction of the PLS2 bundling ability at basic pH appears to be dominated by a weakened affinity of ABD1 to actin. Such domination can be manifested either by hindering the 1 to 2 state transition, if ABD1 is the primary actin-binding domain (Fig. 6B, green panel), or the transition from the state 3’ to state 4, if ABD2 is the primary domain (Fig. 6B, yellow panel). However, the latter case should imply a stronger binding of PLS2 via ABD2, which is not the case. Indeed, the pH dependence of FL PLS2 binding to actin reproduces that of ABD1, as both bind weaker at basic pH (Figs. 1F and 2D), and not that of ABD2, which is stronger at pH 8.0 (Fig. 2E-G). Together, these data favor the “ABD1 binds first” hypothesis (Fig. 6B, green panel), consistent with our previous observations.^27,29^ While compared to ABD2, ABD1 has orders of magnitude lower affinity for actin, its binding in the inhibited state may be favored by a polymorphic nature of this interaction, as judged from the difference revealed from cryo-EM reconstruction of *i)* actin filaments decorated by isolated ABD1^29^ and *ii)* actin bundled by PLS3.^30^

In screening for pH sensor residues, we identified His207 as a likely candidate whose mutagenesis reasonably reproduced the pH-sensitive variations in PLS2 activity. Yet, the introduced constitutively protonated (H207K) and deprotonated (H207Y) residues appear to only partially mimic the pH-sensing mechanisms of WT PLS2. Indeed, H207Y mimicked basic pH effects by moderately inhibiting the interaction between actin and ABD1 at neutral pH (state 1 to 2 transition on Fig. 6B), but it also strengthened the ABD1-ABD2 binding [*i.e.,* inhibited the transition 2 (2’) to 3 (3’)], reproducing the effects of acidic pH on this interaction. H207K caused no influence on the pH sensitivity of ABD1-actin interaction (Fig. 4D) but reduced the affinity of ABD1 to ABD2. Thus, the H207K mutation does not affect the transition from state 1 to 2 (2’) but facilitates the separation of ABDs and the transition from state 2 to 3 or 2’ to 3’, thus favoring the bundling (state 4). Of note, the weakening of the inhibitory ABD1-ABD2 association by H207K likely recapitulates that upon the phosphorylation/mutation of S406.^29^ This similarity should not be surprising given that despite being located in two different ABDs, H207 and S406 are only 4.2 Å apart (Cα-Cα distance) in the AlphaFold2 structure of PLS2 (Fig. S5A,B). While not fully recapitulating the pH sensitivity mechanisms of WT PLS2, the net effects of H207K and H207Y mutations mimic the effects of acidic and basic pH_i_, respectively, and, as such, are valuable as tools for studying the role of pH in PLS2 regulation (Fig. 5).

While searching for residues mediating pH-sensing, we found that H116K strongly inhibits F-actin bundling (Fig. S2A), just as the S117E mutation of the neighboring residue (Fig. S3), suggesting that neither positive nor negative charges are tolerated at this position. Interestingly, the homologous PLS3 residues in a cryo-EM structure of plastin-cross-linked F-actin bundle are disordered and do not obviously contribute to actin binding^30^. In the AlphaFold2-predicted PLS2 structure, these residues form a small antiparallel β-sheet by bonding with residues A102 and I103 (Fig. S5C-E). Whether this region of PLS2 contributes to actin binding directly as the N-terminal residues preceding the *t*-CH domain of β-III-spectrin,^61^ or affect binding indirectly by changing the thermodynamic stability of PLS2 as it was proposed for utrophin,^60^ or by affecting the regulation of PLS2 by RD, remains to be established.

Endogenous expression of PLS2 is restricted to hematopoietic cells,^37^ where it contributes to the formation and stabilization of immune synapses,^66,67^ podosomes^68,69^ and sealing rings, ^70^ but also to the migration of various immune cells.^55,71^ Immune cell migration is known to be inhibited by an acidic environment, a common inflammation hallmark.^72^ One recognized mechanism of this inhibition is a stronger association of integrins with the extracellular matrix at acidic pH.^73^ However, the metabolic drop of the extracellular pH is accompanied by intracellular acidification,^74^ and, therefore, the inhibited migration may also be mediated by pH-sensitive proteins. For PLS2, its stronger bundling capacity in the acidic environment may promote the stability of adhesive structures such as podosomes, which are known to stabilize in response to the acidification of the osteoclast’s cytoplasm.^75,76^ While the potential implications of this regulatory mode are numerous, further studies are needed to clarify the role of PLS2’s pH sensitivity in regulating the activity of immune cells.

Intracellular pH is constitutively elevated in cancer,^3^ where it enhances proliferation, migration, and invasion.^77^ A key requirement for mesenchymal cell migration is the ability to regulate FA turnover^78^. This is achieved by the activation of NHE1, leading to a localized rise in pH_i_ that promotes the disassembly of FAs.^6,8,16^ Both nascent FAs at the leading edge and mature FAs in the cell interior must be regulated to ensure proper migration. PLS2 is ectopically expressed in many cancers,^39–41^ where its presence correlates with their boosted invasive and metastatic properties. Similarly, PLS3 is implicated in cell migration^79,80^ in both healthy and cancerous tissues.^45,81^ Our findings suggest that the pH-sensitivity of PLS2 and PLS3 allows these proteins to respond to increased pH_i_ (*e.g.*, at FAs) by weakening actin bundles and allowing depolymerization factors, such as cofilin and AIP1, to sever and disassemble actin filaments.

PLS2 and PLS3 belong to the plastin/fimbrin family of actin-bundling proteins, which are part of the larger family of *t*-CH actin-binding proteins, the so-called spectrin superfamily.^26^ The ABD shared in each of these proteins has a similar binding footprint on actin^29,30,61,82^ and is hypothesized to share the mechanism of “domain-opening,” which regulates their affinity to F-actin.^29,30,82–85^ With the herein presented example of plastins having pH-dependent F-actin bundling activity and the previously reported pH sensitivity of *D. discoideum* and *H. pulcherrimus* α-actinins^24,25^ it is conceivable that actin binding by other members of the spectrin superfamily may also be sensitive to pH. The hitherto uncharacterized pH sensitivity of these actin-binding proteins may be of considerable importance to disease states such as cancer and neurological disease, both of which display dysregulated intracellular pH.^3,4^

## Methods

### Protein purification and labeling

Skeletal actin was purified from rabbit skeletal muscle acetone powder (Pel-Freeze Biologicals), as previously described^59^. Actin was stored on ice in G-buffer [2 mM Tris-HCl, pH 8.0, 0.2 mM CaCl_2_, 0.2 mM ATP, 5 mM β-mercaptoethanol (β-ME), 0.005% sodium azide] for no longer than one month with dialysis into fresh G-buffer after two weeks of storage.

QuikChange Lightning Multi-Site-Directed Mutagenesis kit (Agilent Technologies) was used to introduce mutations into plastin constructs cloned in-frame with a tobacco etch virus (TEV) protease recognition sequence downstream of the N-terminal 6xHis-tag in pColdI vector^27^. Truncation constructs were created using NEBuilder HiFi DNA assembly (New England Biolabs). All sequences were verified by Sanger DNA sequencing [Genomics shared resource, The Ohio State University Comprehensive Cancer Center (GSR OSUCCC)]. PLS2-S5D was used as a control in all experiments with PLS2 mutants created on the S5D background (*in vitro* properties of PLS2 are not affected by S5D mutation).^27^ Recombinant proteins were expressed in and purified from BL21-CodonPlus(DE3)pLysS *Escherichia coli* (Agilent Technologies) by immobilized metal affinity chromatography (IMAC) as previously described^86^ using HisPur cobalt resin (Thermo Scientific). The 6xHis-tag was removed from all purified proteins by treating them overnight with TEV protease at a 1:20 mole ratio, followed by re-incubation with HisPur cobalt resin to remove the cleaved 6xHis-tag and His-tagged TEV protease. Flow-through fractions containing tagless protein were concentrated and further purified on a Superdex 200 Increase 10/300 GL size-exclusion column (GE Healthcare) equilibrated with PLS buffer [10 mM 4-(2-hydroxyethyl)-1-piperazineethanesulfanoic acid (HEPES), pH 7.0, 30 mM KCl, 2 mM MgCl_2_, 0.5 mM ethylene glycol-bis(β-aminoethyl ether)-N,N,N’,N’-tetraacetic acid (EGTA), 2 mM dithiothreitol (DTT)]. Purified tagless PLS constructs were aliquoted, flash-frozen in liquid nitrogen and stored at −80°C. ABD2 was labeled with fluorescein maleimide (Thermo Fisher Scientific) in PLS buffer devoid of reducing agent at 4°C as previously described.^29^

### Buffer pH determination

The pH values for each buffer condition were carefully controlled via the following procedures. G-buffer was used as the initial solvent, to which all other buffer components (*i.e.,* 10 mM HEPES, 30 mM KCl, 2 mM MgCl_2_, 0.5 mM EGTA) were added from stock solutions to make the PLS buffer with a desired pH measured using a pH Basic meter (Sartorius) with a Fisherbrand Accumet liquid-filled mercury-free pH/ATC electrode (Fisher Scientific). Since the pH of a buffered solution changes in response to ionic strength,^87^ the initial pH parameters of the 1 M HEPES stock solutions were related to the final pH of the PLS buffer solutions by a standard curve of a linear character (Fig. S6). The linear fit yielded Equation 1, which we used to prepare each 1 M HEPES stock solution:

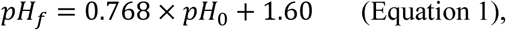

where *pH_f_* is the pH of the final PLS buffer, and *pH_0_* is the pH of the 1 M HEPES stock solution. The 1 M HEPES stock solutions were frozen immediately after preparation and used for the preparation of the final buffers, whose actual pH values were experimentally verified using the pH-meter. Refer to the Supplementary Material (Supplementary Methods section) for additional details.

### Light scattering assays

For each experiment, tagless plastin constructs were cleared by ultracentrifugation at 300,000 g for 30 min at 4°C using an Optima MAX-TL ultracentrifuge (Beckman Coulter). The supernatant was collected, and the remaining soluble protein concentration was determined by absorbance at 280 nm using extinction coefficients determined by ProtParam on the Expasy server.^88^ G-actin in G-buffer was degassed and placed in UV-transparent glass cuvettes; after five minutes, EGTA and MgCl_2_ were added simultaneously to the final concentrations of 0.5 and 0.1 mM, respectively, mixed, and allowed to incubate for another five minutes to switch from Ca^2+^-bound to Mg^2+^-bound actin state. Polymerization was initiated by the addition of the mixture of HEPES, KCl, MgCl_2_, EGTA to the final concentrations specified for the PLS buffer. When bundling was studied, the specified plastin constructs were added at the polymerization initiation time. The final concentrations of actin and plastin were 10 µM and 5.0 µM, respectively. The light scattering intensity change associated with actin polymerization and bundling was measured at 90° to the incident light using a PTI QM-400 fluorometer (Horiba Scientific) with excitation and emission wavelengths set to 350 nm at 25°C. Measurements were taken every ∼10 s, except up to 40 s pauses needed for adding experimental components, which were added to cuvettes in a staggered sequence, with the first and last cuvette initiated at ∼10 min and at ∼25 min, respectively. Since the resulting asynchronous gaps in the data replicates did not allow direct averaging, the traces fit as single, double, or triple exponentials, and the determined parameters were used to extrapolate missing data and fill the gaps. The extrapolated data was averaged and presented in all light scattering curves. Upon completion of the experiment, all samples were pelleted via low-speed co-sedimentation and analyzed by electrophoresis (see below).

### Co-sedimentation assays

Low-speed co-sedimentation assays to monitor F-actin bundling either followed light scattering experiments (as described above) or were conducted as independent experiments. In the latter case, 5 µM actin in G-buffer was incubated with 0.5 mM EGTA and 0.1 mM MgCl_2_ on ice for 5 min, followed by the addition of 1M HEPES of the desired pH, KCl, and MgCl_2_ up to the final concentrations of 10, 30, and 2 mM, respectively, and allowed to polymerize for at least 30 min at 22°C. Plastin constructs of various concentrations were mixed with polymerized actin and incubated overnight at 4°C, followed by further incubation at 22°C for one hour. The reactions were spun at 20,000 g, 25°C, for 20 min. Immediately following the centrifugation, supernatants were separated from pellets and resolved by SDS-PAGE.

During high-speed co-sedimentation, actin was incubated with 0.5 mM EGTA and 0.5 mM MgCl_2_ (higher [Mg^2+^] was used due to the high [actin] >100 µM) on ice for 5 min before the addition of 10 mM HEPES of indicated pH, 30 mM KCl, and 1.5 mM MgCl_2_ to a final concentration of 2 mM. Plastin constructs were mixed with polymerized actin and incubated overnight at 4°C, followed by additional incubation at 22°C for at least one hour. The final concentration of plastin in Figures 1E, 1F, and S1H was 5 µM, while the final concentration of plastin in Figures 2D and 4D was 2 µM. The samples were centrifuged at 300,000 g, 25°C, for 30 min in an Optima MAX-TL ultracentrifuge (Beckman Coulter).

Supernatants were separated from pellets immediately after the spin, pellets were soaked with equivalent volumes of 1x reducing sample buffer at least for 2 hours, collected via vigorous pipetting, and resolved by SDS-PAGE. For both high- and low-speed co-sedimentation, gels were stained with Coomassie Brilliant Blue, and gel band intensities were quantified by densitometry using ImageJ v.2.3 software.^89,90^ The number of repetitions is indicated in the figure legends. Uncropped SDS-PAGE gels are shown in the Appendix A1.

### Stopped-flow kinetics

Time courses of FM-ABD2 binding to or dissociating from F-actin or the indicated RD-ABD1 construct were recorded, and association and dissociation rates were determined by the change in fluorescence anisotropy signal detected by an SX-20 LED stopped-flow spectrometer (Applied Photophysics) at 25°C. The dead time of the instrument is 1 ms. Samples were excited by a 470 nm LED element (Applied Photophysics), and changes in fluorescence anisotropy were measured using two identical 515 nm long-pass colored glass filters (Newport Corporation) in parallel and perpendicular channels. To determine the association rates of FM-ABD2 (50 or 100 nM) with F-actin or the specified RD-ABD1 construct, these proteins were loaded to the instrument syringes in parallel. The unlabeled proteins were present in a range of concentrations at least 10 times higher than that of FM-ABD2 to ensure pseudo-first-order kinetic conditions. Time courses from the association experiments were individually fit to a single exponential equation using Pro-Data SX and Pro-Data Viewer (Applied Photophysics). All *k_obs_* values are the results of at least 8 association time course replicates. Error bars represent the standard deviation (SD) of the mean.

*k_off_* values were directly measured by recording the dissociation of FM-ABD2 from either F-actin or the specified RD-ABD1 construct upon competition with excess of unlabeled ABD2. For dissociation time courses of FM-ABD2 and F-actin, either one of two conditions were used: 500 nM F-actin stabilized with 2 µM phalloidin bound to 50 nM FM-ABD2, or 100 nM F-actin stabilized with 1 µM phalloidin and bound to 25 nM FM-ABD2. For each pH and protein concentration condition, 7.5 and 10 µM unlabeled ABD2 were used to compete with pre-bound FM-ABD2. Both concentrations of unlabeled ABD2 yielded similar rates, confirming that the reassociation of FM-ABD2 was not an issue. These concentrations yielded at least 4 dissociation time courses for each pH condition, single exponential fits of which provided the sought *k_off_* values.

For dissociation time courses with FM-ABD2 and the specified RD-ABD1 construct, 25 or 50 nM FM-ABD2 was bound to 50 or 100 nM RD-ABD1 and competed with 7.5 to 20 µM unlabeled ABD2. These concentrations yielded at least four dissociation time courses for each concentration of ABD2 at each pH. These time courses were individually fit to a single or double exponential, which yielded rate constants that were averaged and taken as the *k_off_*. Both the lower and higher concentrations of ABD2 yielded similar results. All *k_off_* values for FM-ABD2 with F-actin and FM-ABD2 with RD-ABD1 are reported with standard deviation (*n ≥* 4).

To determine *k_on_*, *K_obs_* values measured for association time courses were plotted with the experimentally measured *k_off_* values and fit to a linear line in GraphPad Prism version 10.0.0 for Windows (GraphPad Software, Boston, Massachusetts USA), which was forced to cross the y-axis at the experimentally measured *k_off_*. The slope of this line yielded the *k_on_* and is reported with curve fit error. Representative curves shown in Figures 2F and 4F are averaged, smoothed curves from at least 4 individual time courses, which were fit to a single or double exponential, normalized, and plotted with their best-fit function.

### Fluorescence anisotropy equilibrium binding assays

The change in fluorescence anisotropy of FM-ABD2 at pH 7 and 8 upon binding to either F-actin or RD-ABD1 was measured using an Infinite M1000 Pro plate reader (Tecan US, Inc.) with excitation and emission wavelengths of 470 and 519 nm respectively. Fluorescence anisotropy measurements of F-actin binding were performed after equilibration of 5 nM FM-ABD2 with the indicated concentrations of F-actin stabilized with 1.0 µM phalloidin. RD-ABD1 binding measurements were performed using the indicated concentrations of RD-ABD1, equilibrated with 10 nM FM-ABD2. All equilibrium assays were measured after 60 minutes of incubation at 22°C, followed by an additional observation after 24 h, to confirm that the equilibrium had been reached. All reactions were carried out in PLS buffer of the indicated pH. The data from three technical replicates were normalized and fit to a quadratic isotherm equation:

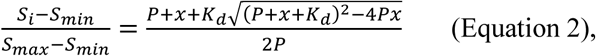

where *S_i_* is the signal for each data point, *S_min_* is the minimum value for the replicate, and *S_max_* is the maximum value from that replicate, *P* is the concentration of labeled FM-ABD2, and *x* is the concentration of either F-actin or RDABD1 construct. K_d_ values obtained from equilibrium measurements are the average of three technical replicates, reported with their standard deviations.

### Cell culture, transfections, microscopy, and pH_i_ clamping and calibration

*Xenopus laevis* XTC fibroblast cells^91^ [obtained from Dr. Watanabe (Kyoto University) and not further authenticated] were cultured in 70% Leibovitz’s L-15 medium (Thermo Fisher Scientific) supplemented with 10% fetal bovine serum, L-glutamine, and penicillin-streptomycin at 23°C and ambient CO_2_. The cells were mycoplasma-negative as determined by PCR.^92^ Transient transfections were performed using Lipofectamine 3000 (Thermo Fisher Scientific) and human PLS2 constructs N-terminally fused to mEmerald.^29^ To introduce the desired mutations, mutagenesis was performed using QuikChange Site-Directed Mutagenesis kit (Agilent Technologies). mCherry-β-actin (Addgene #54967, RRID:Addgene_54967) was a gift from Michael Davidson.^93^ mCherry-SEpHluorin (Addgene plasmid #32001; RRID:Addgene_32001) was a gift from Sergio Grinstein.^94^ Transfected cells were plated on polylysine-coated coverslips (Neuvitro Corporation, Vancouver, WA) in Attofluor chambers (Thermo Fisher Scientific, Waltham, MA) in serum-free L-15 medium and imaged 30 min post-plating. Micrographs were obtained using a Nikon Eclipse Ti-E inverted microscope (Nikon Instruments Inc., Melville, NY) equipped with a perfect focus system, Nikon CFI Plan Apochromat λ 100x oil objective (NA 1.45), and an iXon Ultra 897 EMCCD camera (Andor Technology, Belfast, UK) using NIS Elements-AR v.4.3 software (Nikon Instruments Inc., Melville, NY).

For pH_i_ calibration curve (Fig. S4A,B), cells transiently transfected with mCherry-SEpHluorin were sequentially incubated and imaged in nigericin buffers (10 µM nigericin, 25 mM HEPES, 105 mM KCl, 1 mM MgCl_2_) adjusted to a desired pH using KOH to satisfy the Na^+^-free buffer requirement for equilibration of intracellular and extracellular pH via nigericin clamping.^64,65^ Cells were fragmented using Threshold in Fiji/ImageJ2.^89^ The ratios of the background-corrected fluorescence intensities of SEpHluorin (pH sensor) to mCherry (pH-insensitive) were plotted against the corresponding pH value. Data are presented as means ± SD; individual numbers of analyzed cells (n) for each pH condition are given in Figure S4B legend.

Time-lapse imaging (Fig. 5; Videos S1-4) was performed on cells transiently co-transfected with mCherry-β-actin and various mEmerald-PLS2 constructs at intervals of 1 min for 10 min under the pre-clamping conditions. Then, while on the microscope stage, cell medium was replaced with nigericin buffer of a desired pH, and imaging was continued for additional 25 min before changing to a different pH buffer with an additional 25-min imaging. Kymographs and line plot profiles were obtained using KimographBuilder and Plot Profile in Fiji/ImageJ2. In additional experiments, cells were cycled through several changes of nigericin buffers with different pH, and random transfected cells were imaged at each pH_i_ condition tested (Fig. S4C,D).

### AlphaFold2-generated images

AlphaFold2^95,96^ was used to generate the structure of PLS2 (Fig. S5).

### Statistical analysis

*P* values were calculated using Student’s t-test in Microsoft Excel for Microsoft 365 MSO version 2307 (Microsoft Corporation) when comparing only two groups (Figs. 1E, 2C and D, 4C). All other multiple comparisons were done using analysis of variance (ANOVA) with a Turkey’s post-hoc test in Origin 2023 v.10.0 or GraphPad PRISM v. 10.0.0 for Windows. Individual *P* values for the experimental data from Figure 1D are presented in Tables S1 and S2. Individual *P* values for the experimental data from Figure 4D are presented in Table S4. Results were considered significant if the associated *P* value was less than 0.05. The definition of error bars and number of repetitions are described in figure legends. Graphical data is provided in the Appendix A2.

## CRediT authorship contribution statement

All authors: investigation; LAR, EK: formal analysis; EK, LAR, DSK: visualization; LAR, DSK, EK: writing – original draft; LAR, EK, DSK: writing – review & editing; DSK: supervision, funding acquisition.

## Data availability

The authors declare that the data supporting the findings of this study are available within the article and its Supplementary Material and Appendices files.

## Abbreviations

ABD: actin-binding domain
AIP1: actin-interacting protein 1
CBM: calmodulin-binding motif
CH domain: calponin-homology domain
F-actin: filamentous actin
FA: focal adhesion
FAK: focal adhesion kinase
FL: full-length
NHE1: Na^+^/H^+^ exchanger 1
pH_i_: intracellular pH
PLS: plastin
RD: regulatory domain
*t*-CH domain: tandem calponin-homology domain
WT: wild type

## References

1. Casey, J. R., Grinstein, S. & Orlowski, J. Sensors and regulators of intracellular pH. Nat. Rev. Mol. Cell Biol. 11, 50–61 (2010).

2. Spear, J. S. & White, K. A. Single-cell intracellular pH dynamics regulate the cell cycle by timing the G1 exit and G2 transition. J. Cell Sci. 136, 2021.06.04.447151 (2023).

3. Czowski, B. J., Romero-Moreno, R., Trull, K. J. & White, K. A. Cancer and pH dynamics: Transcriptional regulation, proteostasis, and the need for new molecular tools. Cancers (Basel). 12, 1–19 (2020).

4. Majdi, A. et al. Permissive role of cytosolic pH acidification in neurodegeneration: A closer look at its causes and consequences. J. Neurosci. Res. 94, 879–887 (2016).

5. Stock, C. & Schwab, A. Role of the Na + /H + exchanger NHE1 in cell migration. Acta Physiol. 187, 149–157 (2006).

6. Ludwig, F. T., Schwab, A., Stock, C. & Al, L. E. T. The Na+ / H+ - Exchanger (NHE1) Generates pH Nanodomains at Focal Adhesions. J. Cell. Physiol. 228, 1351–1358 (2012).

7. Denker, S. P. & Barber, D. L. Cell migration requires both ion translocation and cytoskeletal anchoring by the Na-H exchanger NHE1. J. Cell Biol. 159, 1087–1096 (2002).

8. Choi, C., Webb, B. A., Chimenti, M. S., Jacobson, M. P. & Barber, D. L. pH sensing by FAK-His58 regulates focal adhesion remodeling. J. Cell Biol. 202, 849–859 (2013).

9. Schwab, A., Fabian, A., Hanley, P. J. & Stock, C. Role of Ion Channels and Transporters in Cell Migration. Physiol. Rev. 92, 1865–1913 (2012).

10. Cardone, R. A., Casavola, V. & Reshkin, S. J. The role of disturbed pH dynamics and the NA+/H+ exchanger in metastasis. Nat. Rev. Cancer 5, 786–795 (2005).

11. Kruse, C. R. et al. The effect of pH on cell viability, cell migration, cell proliferation, wound closure, and wound reepithelialization: In vitro and in vivo study. Wound Repair Regen. 25, 260– 269 (2017).

12. Martin, C., Pedersen, S. F., Schwab, A. & Stock, C. Intracellular pH gradients in migrating cells. Am. J. Physiol. - Cell Physiol. 300, C490–C495 (2011).

13. Frantz, C., Karydis, A., Nalbant, P., Hahn, K. M. & Barber, D. L. Positive feedback between Cdc42 activity and H+ efflux by the Na-H exchanger NHE1 for polarity of migrating cells. J. Cell Biol. 179, 403–410 (2007).

14. Pope, B. J., Zierler-Gould, K. M., Kühne, R., Weeds, A. G. & Ball, L. J. Solution structure of human cofilin: Actin binding, pH sensitivity, and relationship to actin-depolymerizing factor. J. Biol. Chem. 279, 4840–4848 (2004).

15. Frantz, C. et al. Cofilin is a pH sensor for actin free barbed end formation: role of phosphoinositide binding. J. Cell Biol. 183, 865–879 (2008).

16. Srivastava, J. et al. Structural model and functional significance of pH-dependent talin–actin binding for focal adhesion remodeling. Proc. Natl. Acad. Sci. 105, 14436–14441 (2008).

17. Magalhaes, M. A. O. et al. Cortactin phosphorylation regulates cell invasion through a pH-dependent pathway. J. Cell Biol. 195, 903–920 (2011).

18. Nomura, K., Hayakawa, K., Tatsumi, H. & Ono, S. Actin-interacting protein 1 promotes disassembly of actin-depolymerizing factor/cofilin-bound Actin filaments in a pH-dependent manner. J. Biol. Chem. 291, 5146–5156 (2016).

19. Hall, A. Rho GTPases and the control of cell behaviour. Biochem. Soc. Trans. 33, 891–895 (2005).

20. Hawkins, M., Pope, B., Maciver, S. K. & Weeds, A. G. Human Actin Depolymerizing Factor Mediates Actin Filaments pH-Sensitive Destruction of Actin Filaments. Biochemistry 9985–9993 (1993).

21. Chen, H. et al. In Vitro Activity Differences between Proteins of the ADF / Cofilin Family Define Two Distinct Subgroups. Biochemistry 7127–7142 (2004).

22. Wioland, H., Jegou, A. & Romet-lemonne, G. Quantitative Variations with pH of Actin Depolymerizing Factor/ Cofilin’s Multiple Actions on Actin Filaments. Biochemistry 40–47 (2019) doi:10.1021/acs.biochem.8b01001.

23. DesMarais, V., Ghosh, M., Eddy, R. & Condeelis, J. Cofilin takes the lead. J. Cell Sci. 118, 19–26 (2005).

24. Condeelis, J. & Vahey, M. A Calcium- and pH-regulated Protein from Dictyostelium discoideum That Cross-links Actin Filaments. J. Cell Biol. 94, 466–471 (1982).

25. Mabuchi, I. et al. Alpha-Actinin from Sea Urchin Eggs: Biochemical Properties, Interaction with Actin, and Distribution in the Cell during Fertilization and Cleavage. J. Cell Biol. 100, 375–383 (1985).

26. Liem, R. K. H. Cytoskeletal integrators: The spectrin superfamily. Cold Spring Harb. Perspect. Biol. 8, 1–10 (2016).

27. Schwebach, C. L., Agrawal, R., Lindert, S., Kudryashova, E. & Kudryashov, D. S. The Roles of Actin-Binding Domains 1 and 2 in the Calcium-Dependent Regulation of Actin Filament Bundling by Human Plastins. J. Mol. Biol. 429, 2490–2508 (2017).

28. Klein, M. G. et al. Structure of the actin crosslinking core of fimbrin. Structure 12, 999–1013 (2004).

29. Schwebach, C. L. et al. Allosteric regulation controls actin-bundling properties of human plastins. Nat. Struct. Mol. Biol. 29, 519–528 (2022).

30. Mei, L. et al. Structural mechanism for bidirectional actin cross-linking by T-plastin. Proc. Natl. Acad. Sci. U. S. A. 119, 1–11 (2022).

31. Blanchoin, L., Boujemaa-Paterski, R., Sykes, C. & Plastino, J. Actin dynamics, architecture, and mechanics in cell motility. Physiol. Rev. 94, 235–263 (2014).

32. Pollard, T. D. Actin and actin-binding proteins. Cold Spring Harb. Perspect. Biol. 8, (2016).

33. Lin, C. S. S. et al. Human plastin genes. Comparative gene structure, chromosome location, and differential expression in normal and neoplastic cells. J. Biol. Chem. 268, 2781–2792 (1993).

34. Shinomiya, H. Plastin Family of Actin-Bundling Proteins: Its Functions in Leukocytes, Neurons, Intestines, and Cancer. Int. J. Cell Biol. 2012, 1–8 (2012).

35. Lin, C. et al. Identification of I-Plastin, a Human Fimbrin Isoform Expressed in Intestine and Kidney. Mol. Cell. Biol. 14, 2457–2467 (1994).

36. Diaz-Horta, O. et al. Novel variant p.E269K confirms causative role of PLS1 mutations in autosomal dominant hearing loss. Clin. Genet. 96, 575–578 (2019).

37. Morley, S. C. The Actin-Bundling Protein L-Plastin: A Critical Regulator of Immune Cell Function. Int. J. Cell Biol. 2012, 1–10 (2012).

38. Wolff, L. et al. Plastin 3 in health and disease: a matter of balance. Cell. Mol. Life Sci. (2021) doi:10.1007/s00018-021-03843-5.

39. Park, T., Chen, Z. P. & Leavitt, J. Activation of the Leukocyte Plastin Gene Occurs in Most Human Cancer Cells. Cancer Res. 54, 1775–1781 (1994).

40. Foran, E., McWilliam, P., Kelleher, D., Croke, D. T. & Long, A. The leukocyte protein L-plastin induces proliferation, invasion and loss of E-cadherin expression in colon cancer cells. Int. J. Cancer 118, 2098–2104 (2006).

41. Riplinger, S. M. et al. Metastasis of prostate cancer and melanoma cells in a preclinical in vivo mouse model is enhanced by L-plastin expression and phosphorylation. Mol. Cancer 13, 1–12 (2014).

42. Machado, R. A. C. et al. L-plastin Ser5 phosphorylation is modulated by the PI3K/SGK pathway and promotes breast cancer cell invasiveness. Cell Commun. Signal. 19, 1–22 (2021).

43. Ma, Y. et al. Plastin 3 down-regulation augments the sensitivity of MDA-MB-231 cells to paclitaxel via the p38 MAPK signalling pathway. *Artif. Cells, Nanomedicine*, Biotechnol. 47, 684– 694 (2019).

44. Bosseler, M. et al. Inhibition of HIF1α-Dependent Upregulation of Phospho-l-Plastin Resensitizes Multiple Myeloma Cells to Frontline Therapy. Int. J. Mol. Sci. 19, 1551 (2018).

45. Xin, Z. et al. PLS3 predicts poor prognosis in pancreatic cancer and promotes cancer cell proliferation via PI3K/AKT signaling. J. Cell. Physiol. 235, 8416–8423 (2020).

46. Wang, D. et al. PLS3 promotes papillary thyroid carcinoma progression by activating the Notch signaling pathway. Environ. Toxicol. 39, 539–550 (2024).

47. Janji, B. et al. Phosphorylation on Ser5 increases the F-actin-binding activity of L-plastin and promotes its targeting to sites of actin assembly in cells. J. Cell Sci. 119, 1947–1960 (2006).

48. Schwebach, C. L. et al. Osteogenesis imperfecta mutations in plastin 3 lead to impaired calcium regulation of actin bundling. Bone Res. (2020) doi:10.1038/s41413-020-0095-2.

49. Ishida, H., Jensen, K. V., Woodman, A. G., Hyndman, M. E. & Vogel, H. J. The Calcium-Dependent Switch Helix of L-Plastin Regulates Actin Bundling. Sci. Rep. 7, 1–12 (2017).

50. Huttlin, E. L. et al. A Tissue-Specific Atlas of Mouse Protein Phosphorylation and Expression. Cell 143, 1174–1189 (2010).

51. Klammer, M. et al. Phosphosignature predicts dasatinib response in non-small cell lung cancer. Mol. Cell. Proteomics 11, 651–668 (2012).

52. Mertins, P. et al. Proteogenomics connects somatic mutations to signalling in breast cancer. Nature 534, 55–62 (2016).

53. Mertins, P. et al. Ischemia in Tumors Induces Early and Sustained Phosphorylation Changes in Stress Kinase Pathways but Does Not Affect Global Protein Levels. Mol. Cell. Proteomics 13, 1690–1704 (2014).

54. Zhou, H. et al. Toward a comprehensive characterization of a human cancer cell phosphoproteome. J. Proteome Res. 12, 260–271 (2013).

55. Freeley, M. et al. L-Plastin Regulates Polarization and Migration in Chemokine-Stimulated Human T Lymphocytes. J. Immunol. 188, 6357–6370 (2012).

56. Namba, Y., Ito, M., Zu, Y., Shigesada, K. & Maruyama, K. Human T cell L-plastin bundles actin filaments in a calcium dependent manner. J. Biochem. 112, 503–507 (1992).

57. Ellis, K. J. & Morrison, J. F. Buffers of Constant Ionic Strength for Studying pH-Dependent Processes. Biochemistry 20, 1805 (1981).

58. Wang, F., Sampogna, R. V. & Ware, B. R. pH dependence of actin self-assembly. Biophys. J. 55, 293–298 (1989).

59. Pardee, J. D. & Spudich, J. A. Purification of Muscle Actin. Methods Cell Biol. 24, 271–289 (1982).

60. Singh, S. M., Bandi, S. & Mallela, K. M. G. The N-Terminal Flanking Region Modulates the Actin Binding Affinity of the Utrophin Tandem Calponin-Homology Domain. Biochemistry 56, 2627–2636 (2017).

61. Avery, A. W. et al. Structural basis for high-affinity actin binding revealed by a β-III-spectrin SCA5 missense mutation. Nat. Commun. 8, 1–7 (2017).

62. Jarmoskaite, I., Alsadhan, I., Vaidyanathan, P. P. & Herschlag, D. How to measure and evaluate binding affinities. Elife 9, 1–34 (2020).

63. Edgcomb, S. P. & Murphy, K. P. Variability in the pKa of Histidine Side-Chains Correlates With Burial Within Proteins. Proteins Struct. Funct. Bioinforma. 49, 1–6 (2002).

64. Thomas, J. A., Buchsbaum, R. N., Zimniak, A. & Racker, E. Intracellular pH measurements in Ehrlich ascites tumor cells utilizing spectroscopic probes generated in situ. Biochemistry 18, 2210–8 (1979).

65. Grillo-Hill, B. K., Webb, B. A. & Barber, D. L. Ratiometric imaging of pH probes. Methods Cell Biol. 123, 429–48 (2014).

66. Wabnitz, G. H. et al. Sustained LFA-1 cluster formation in the immune synapse requires the combined activities of L-plastin and calmodulin. Eur. J. Immunol. 40, 2437–2449 (2010).

67. Wang, C. et al. Actin-Bundling Protein L-Plastin Regulates T Cell Activation. J. Immunol. 185, 7487–7497 (2010).

68. Zhou, J. Y. et al. L-Plastin promotes podosome longevity and supports macrophage motility. Mol Immunol October, 79–88 (2016).

69. Linehan, J. B., Lucas Zepeda, J., Mitchell, T. A. & LeClair, E. E. Follow that cell: Leukocyte migration in L-plastin mutant zebrafish. Cytoskeleton 79, 26–37 (2022).

70. Chellaiah, M. A. et al. L-Plastin deficiency produces increased trabecular bone due to attenuation of sealing ring formation and osteoclast dysfunction. Bone Res. 8, (2020).

71. Todd, E. M. et al. The Actin-Bundling Protein L-Plastin Is Essential for Marginal Zone B Cell Development. J. Immunol. 187, 3015–3025 (2011).

72. Lardner, A. The effects of extracellular pH on immune function. J. Leukoc. Biol. 69, 522–530 (2001).

73. Paradise, R. K., Lauffenburger, D. A. & Van Vliet, K. J. Acidic Extracellular pH Promotes Activation of Integrin αvβ3. PLoS One 6, e15746 (2011).

74. Salameh, A. I., Ruffin, V. A. & Boron, W. F. Effects of metabolic acidosis on intracellular pH responses in multiple cell types. Am J Physiol Regul Integr Comp Physiol 307, 1413–1427 (2014).

75. Teti, A. et al. Regulation of podosomes by intracellular pH in avian osteoclasts. Boll. Soc. Ital. Biol. Sper. 65, 597–601 (1989).

76. Teti, A. et al. Extracellular protons acidify osteoclasts, reduce cytosolic calcium, and promote expression of cell-matrix attachment structures. J. Clin. Invest. 84, 773–780 (1989).

77. Webb, B. A., Chimenti, M., Jacobson, M. P. & Barber, D. L. Dysregulated pH: a perfect storm for cancer progression. Nat. Publ. Gr. 11, 671–677 (2011).

78. O’Neill, G. M. The coordination between actin filaments and adhesion in mesenchymal migration. Cell Adhes. Migr. 3, 2–5 (2009).

79. Brun, C. et al. T-plastin expression downstream to the calcineurin/NFAT pathway is involved in keratinocyte migration. PLoS One 9, 1–8 (2014).

80. Garbett, D. et al. T-Plastin reinforces membrane protrusions to bridge matrix gaps during cell migration. Nat. Commun. 11, (2020).

81. Park, S. Y. et al. T-plastin contributes to epithelial-mesenchymal transition in human lung cancer cells through FAK/AKT/Slug axis signaling pathway. BMB Rep. 57, 305–310 (2024).

82. Galkin, V. E., Orlova, A., Salmazo, A., Djinovic-Carugo, K. & Egelman, E. H. Opening of tandem calponin homology domains regulates their affinity for F-actin. Nat. Struct. Mol. Biol. 17, 614–616 (2010).

83. Lin, A. Y., Prochniewicz, E., James, Z. M., Svensson, B. & Thomas, D. D. Large-scale opening of utrophin’s tandem calponin homology (CH) domains upon actin binding by an induced-fit mechanism. Proc. Natl. Acad. Sci. U. S. A. 108, 12729–12733 (2011).

84. Fealey, M. E. et al. Dynamics of Dystrophin’s Actin-Binding Domain. Biophys. J. 115, 445–454 (2018).

85. Harris, A. R. et al. Steric regulation of tandem calponin homology domain actin-binding affinity. Mol. Biol. Cell 30, 3112–3122 (2019).

86. Dong, S. et al. Photorhabdus luminescens TccC3 Toxin Targets the Dynamic Population of F-Actin and Impairs Cell Cortex Integrity. Int. J. Mol. Sci. 23, 7026 (2022).

87. Voinescu, A. E. et al. Similarity of salt influences on the pH of buffers, polyelectrolytes, and proteins. J. Phys. Chem. B 110, 8870–8876 (2006).

88. Wilkins, M. R. et al. Protein identification and analysis tools in the ExPASy server. Methods Mol. Biol. 112, 531–52 (1999).

89. Schindelin, J., et al. Fiji: an open-source platform for biological-image analysis. Nat. Methods 9, 676–682 (2012).

90. Schindelin, J., Rueden, C. T., Hiner, M. C. & Eliceiri, K. W. The ImageJ ecosystem: An open platform for biomedical image analysis. Mol. Reprod. Dev. 82, 518–529 (2015).

91. Watanabe, N. Fluorescence single-molecule imaging of actin turnover and regulatory mechanisms. Methods in Enzymology vol. 505 (Elsevier Inc., 2012).

92. Uphoff, C. C. & Drexler, H. G. Detection of Mycoplasma Contamination in Cell Cultures. Curr. Protoc. Mol. Biol. 106, (2014).

93. Rizzo, M. A., Davidson, M. W. & Piston, D. W. Fluorescent protein tracking and detection: fluorescent protein structure and color variants. Cold Spring Harb. Protoc. 2009, pdb.top63 (2009).

94. Koivusalo, M. et al. Amiloride inhibits macropinocytosis by lowering submembranous pH and preventing Rac1 and Cdc42 signaling. J. Cell Biol. 188, 547–563 (2010).

95. Jumper, J. et al. Highly accurate protein structure prediction with AlphaFold. Nature 596, 583– 589 (2021).

96. Varadi, M. et al. AlphaFold Protein Structure Database: Massively expanding the structural coverage of protein-sequence space with high-accuracy models. Nucleic Acids Res. 50, D439– D444 (2022).

