## Supplementary materials for "Human plastins are novel cytoskeletal pH sensors with a reduced F-actin bundling capacity at basic pH"

This file includes:

Supplementary Methods

Supplementary Figures (S1-S6)

Supplementary Tables (S1-S5)

Supplementary Video Captions (S1-S4)

### Supplementary Methods

*Calculation of ionic strength equivalent for titration of HEPES buffer to pH 8.0*

To calculate the ionic strength contributed by the addition of KOH (and the resulting deprotonated HEPES buffer) required to adjust the pH of HEPES buffer from pH 7.0 to pH 8.0, we followed the method described by Ellis & Morrsion.[1] At pH 7.0, 10 mM HEPES ($pK_a^*$ = 7.39[1]) buffer solution will be composed of 7.1 mM protonated HEPES (HA, uncharged) and 2.9 mM deprotonated HEPES ($A^-$, negatively charged). Since the concentration of $H^+$ and $OH^-$ are negligible,[1] the total charge of the buffered solution is equal to the amount of strong acid or base required to arrive at the specified pH. The total charge of a pH 7.0 10 mM HEPES is −2.9 mM, which yields an ionic strength of 2.9 mM. Using this same method, the total charge and ionic strength of pH 8.0 10 mM HEPES is −8.8 mM and 8.8 mM, respectively. The difference in ionic strength between these two buffers is 5.9 mM. Therefore, we added 5.9 mM KCl (total of 36 mM KCl) to pH 7.0 PLS buffer to bring it to identical ionic strength with pH 8.0 PLS buffer. Conversely, in a separate experiment, we decreased the concentration of KCl in pH 8.0 PLS buffer to 24.1 mM, to cause the ionic strength of the pH 8.0 buffer to become equal to that of the pH 7.0 buffer.

References

1. Ellis, K. J. & Morrison, J. F. Buffers of Constant Ionic Strength for Studying pH-Dependent Processes. *Biochemistry* **20**, 1805 (1981).

### Supplementary Figures

#### Figure S1

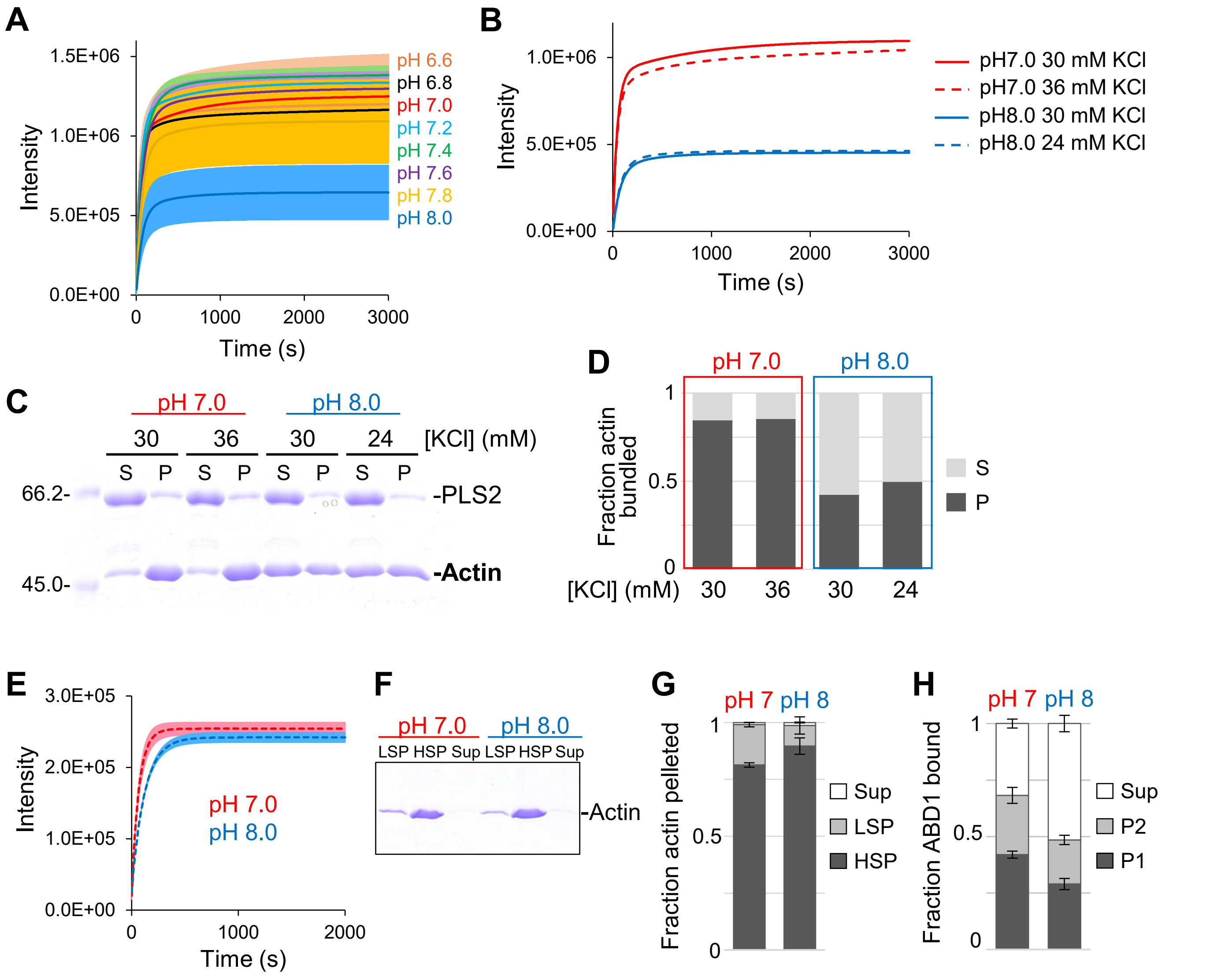


**Figure S1. Characterization of pH-dependence of PLS2 and F-actin.** (A) Light scattering traces of F-actin bundling by PLS2 at various pH values (some are the same as shown on Fig. 1B). Solid colored lines represent averaged extrapolated data, colored areas represent SD of the mean ($n \geq 3$). (B) Light scattering traces of F-actin bundling by PLS2 at pH 7.0 and 8.0 under different salt conditions (to account for an additional ionic strength equivalent (6 mM KCl) during buffer pH adjustments (see Supplementary Methods). (C) Representative 10% SDS-PAGE gel of supernatant (S) and pellet (P) fractions from low-speed co-sedimentation of actin bundles formed by PLS2 in different pH and salt conditions. (D) Fractions of actin in the pellet (P) and supernatant (S) after low-speed co-sedimentation in the presence of PLS2 in different pH and salt conditions quantified from (C). (E) Light scattering traces of polymerization of F-actin alone at pH 7.0 and pH 8.0: a full-scale graph of the "Actin alone" traces, which are shown in Fig. 1B. Solid lines represent averaged extrapolated data, colored areas represent SD of the mean ($n = 2$). (F) Representative 10% SDS-PAGE gel of actin fractions from the light scattering experiments (shown in E): LSP, pelleted at low speed; HSP, pelleted at high speed; Sup, supernatant after the high-speed centrifugation. (G) Quantitation of the gel (F). Error bars represent SD of the mean ($n = 2$). (H) High-speed co-sedimentation experiments of ABD1/F-actin binding: P1 represents fraction of ABD1 present after initial pelleting with 50 μM F-actin; the resulting supernatant was supplemented with another 50 μM F-actin, and pelleted again, yielding another pellet (P2) and a final supernatant (Sup). Error bars represent SD of the mean ($n = 3$).

**Figure S2**

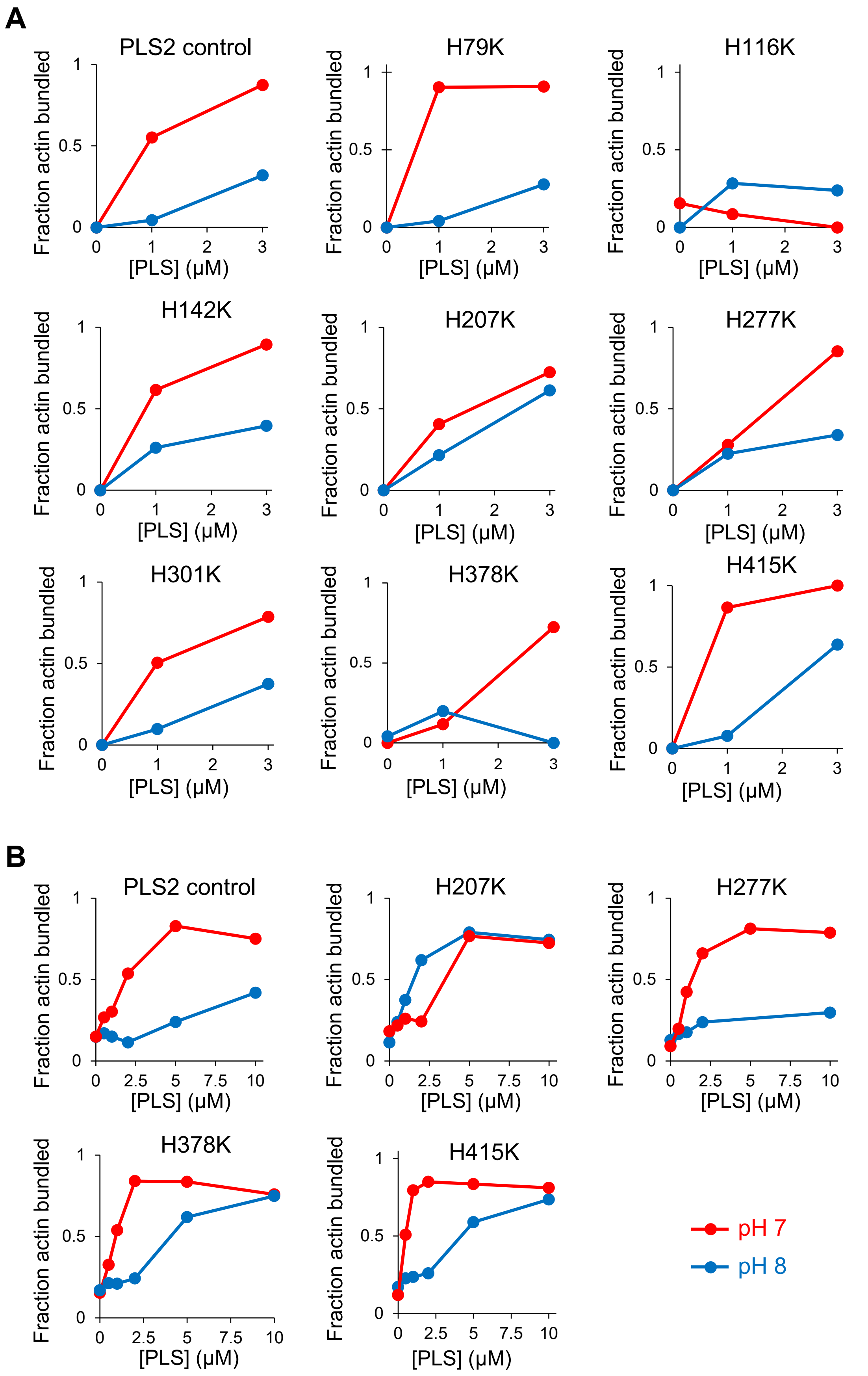


**Figure S2. Screening for pH-dependence of F-actin bundling by PLS2 variants with His-to-Lys substitutions.** (A) Low-speed co-sedimentation of F-actin with indicated PLS2 variants at pH 7.0 (red) and pH 8.0 (blue). (B) Additional low-speed actin co-sedimentation experiments of selected PLS2 constructs with additional titration points.

### Figure S3

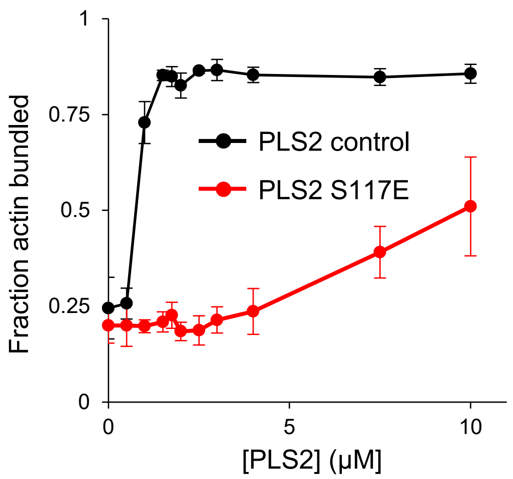


**Figure S3. Effect of S117E mutation on F-actin bundling by PLS2 using low-speed actin co-sedimentation.** Data are presented as mean ± SD; n = 3 (except for PLS2 control at 2μM, S117E at 0.5, 1.75, and 7.5 μM, where n = 2).

**Figure S4**

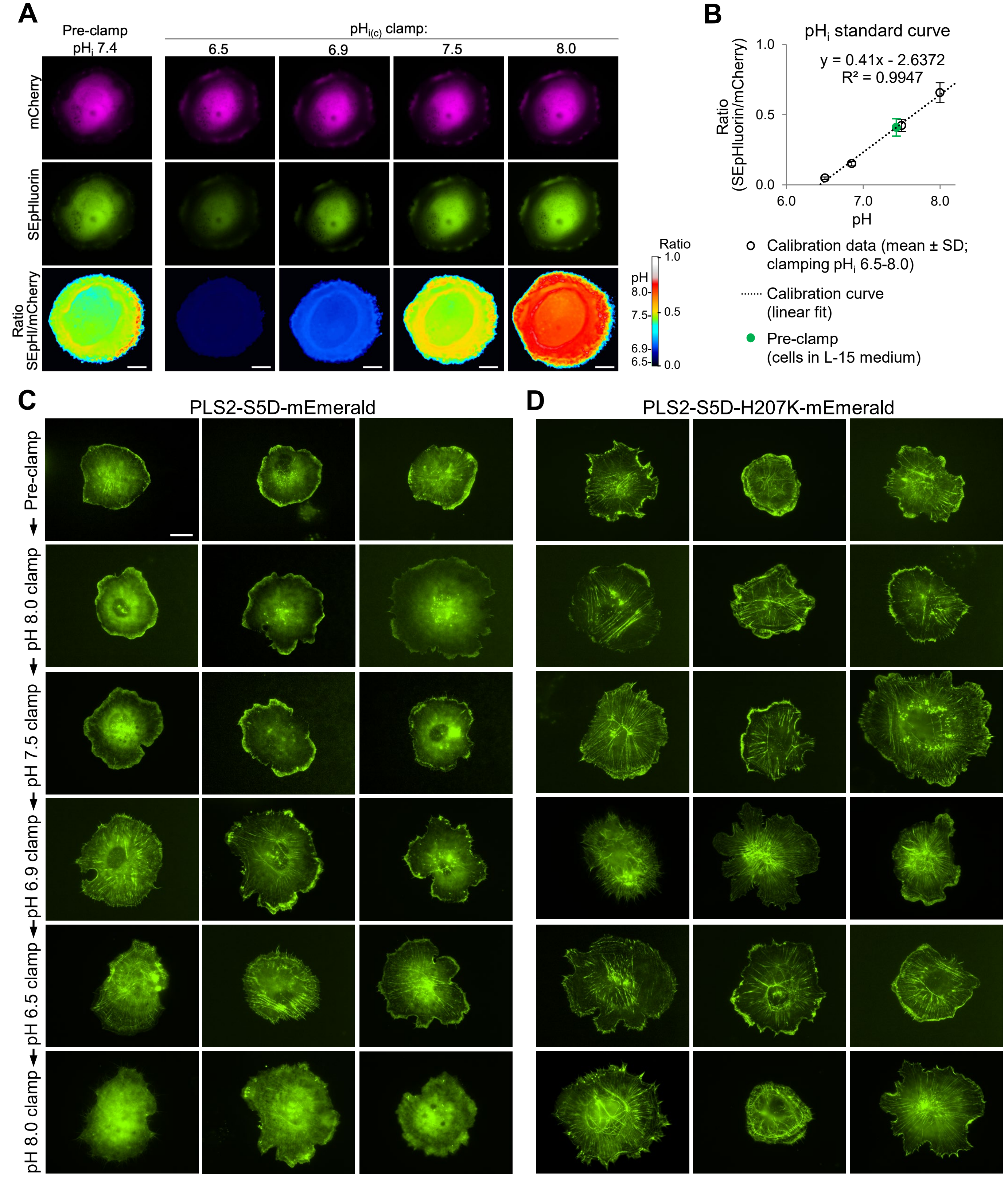


**Figure S4. Effects of intracellular pH$_i$ on PLS2 distribution.** (A) Representative ratiometric images of cells expressing mCherry-SEpHluorin before and after clamping in nigericin buffers of indicated pH. Ratio of background-corrected fluorescence signals in green versus red channels is presented using royal LUT after applying median filter (radius 15 px). Scale bars are 20 µm. (B) pH$_i$ standard curve derived from the ratiometric imaging. Data are presented as mean ± SD; number of analyzed cells (n): pre-clamping, n=24; pH$_i$ 6.5, n=29; pH$_i$ 6.9, n=28; pH$_i$ 7.5, n=29; pH$_i$ 8.0, n=27. (C, D) Representative images of cells expressing mEmerald-tagged PLS2 constructs (control S5D or S5D-H207K) under different pH$_i$ conditions. Cells were clamped at different pH sequentially (as shown by arrows on the left side). Three individual random cells are shown for each pH$_i$ condition for each construct. Scale bar is 20 µm (identical for all images in C and D).

**Figure S5**

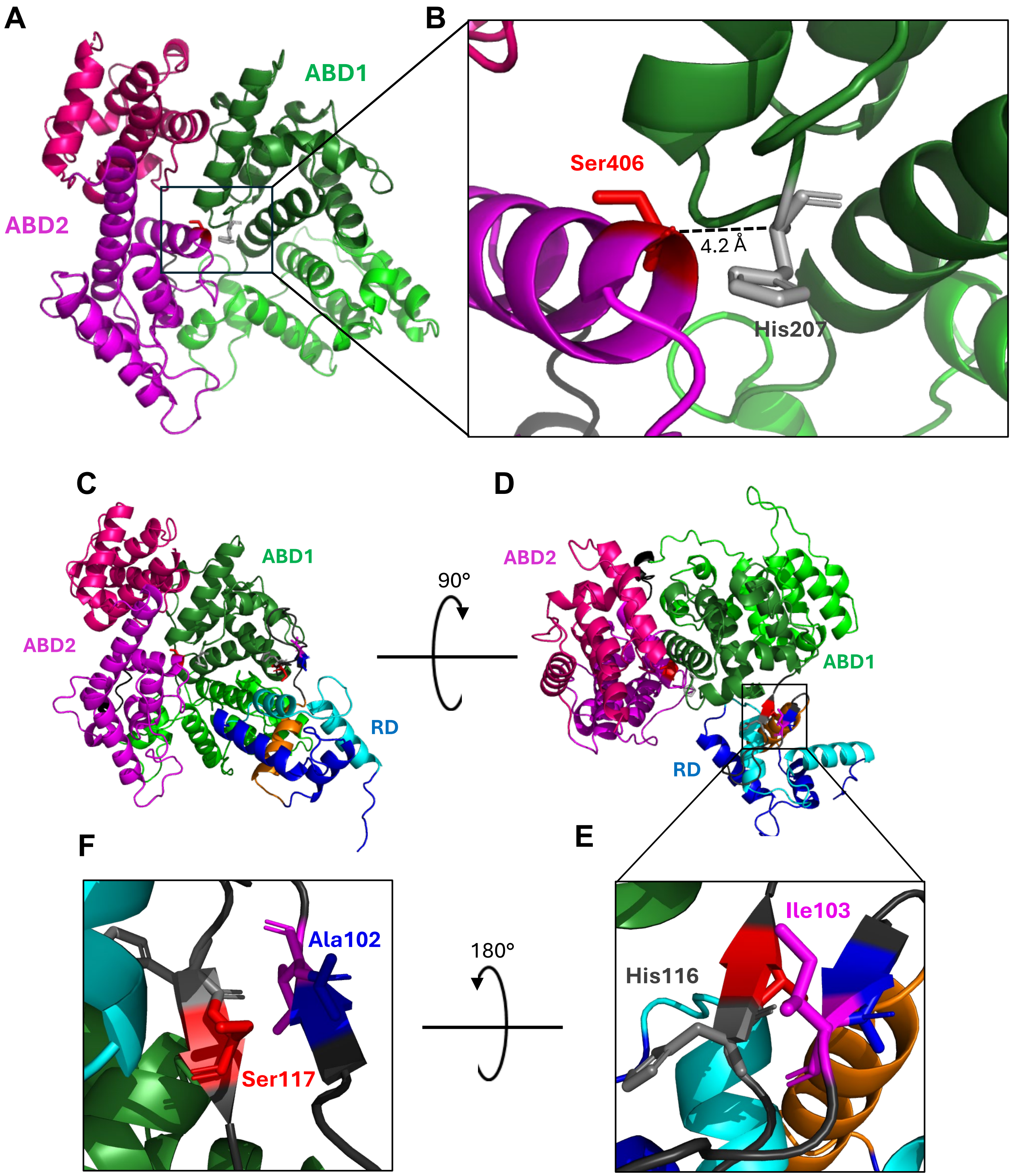


**Figure S5. Relative location of selected residues in tertiary structure of PLS2 as predicted by AlphaFold2**. (A,B) Actin-binding core of PLS2 (RD is not shown). Domains are indicated by color: dark green, CH1; green, CH2; magenta, CH3; pink, CH4. (B) is zoomed image of (A) to show location of Ser406 (red sticks) and His207 (grey sticks). Distance between the α-carbon atoms of Ser406 and His207 is indicated as black dashed line. (C-F) Full-length PLS2. Domains colored as in A, with additional domains indicated by color: light blue, EF-1; blue, EF-2; orange, CBM; dark grey, linker region between CBM and ABD1 (a.a. 97 to 120). Selected residues located in the linker between ABD1 and CBM are shown as sticks: Ser117 (red), His116 (grey), Ala102 (blue), and Ile103 (magenta). (D) Full-length PLS2 as in (C) rotated 90° about the y-axis. (E) A zoomed image of (D) to show relative location of His116 and its H-bonding partner Ile103 in an anti-parallel β-sheet. (F) image of (E) rotated 180° about the y-axis to show relative location of Ser117 and its H-bonding partner Ala102 in an antiparallel β-sheet. Notice that the RD position relative to the core is unlikely to be predicted correctly.

#### Figure S6

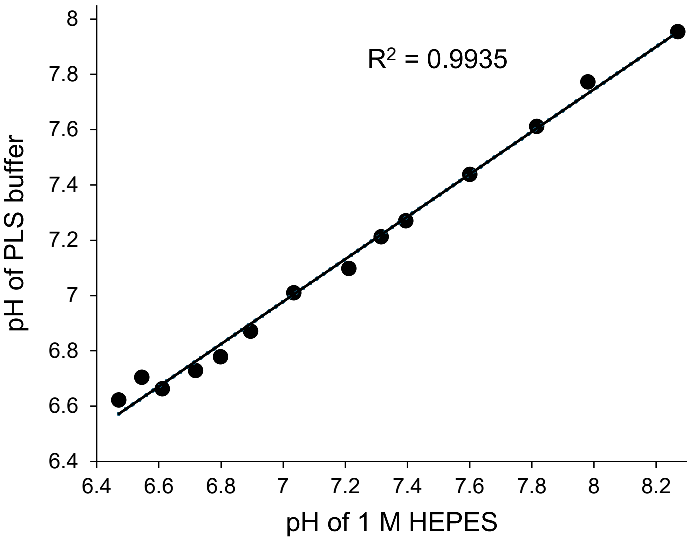


**Figure S6. Standard curve for PLS buffer pH estimation.** Final pH values of assembled PLS buffers are plotted against the pH of 1 M HEPES stock solution used to produce each of them. Dots represent data; line is a linear fit. Related to Methods (*Buffer pH determination* section).

### Supplementary Tables

#### Table S1

**Analysis of variance with Turkey's post-hoc test on the data from light scattering of F-actin in the presence of PLS2 with varying pH.** Final light scattering intensity at 3000 s for each pH condition compared to each other pH condition; n.s. is non-significant.
Related to Figure 1D.

| pH condition | Δ mean | SE of mean | q Value | Probability | Alpha | Significance | Lower control limit | Upper control limit |
|---|---|---|---|---|---|---|---|---|
| pH 6.8 vs pH 6.6 | -35260.57719 | 170153.2098 | 0.29307 | 1 | 0.05 | n.s. | -615701.5249 | 545180.3705 |
| pH 7.0 vs pH 6.6 | 49031.26582 | 159163.7536 | 0.43566 | 0.99998 | 0.05 | n.s. | -493921.5241 | 591984.0557 |
| pH 7.0 vs pH 6.8 | 84291.84301 | 159163.7536 | 0.74896 | 0.99929 | 0.05 | n.s. | -458660.9469 | 627244.6329 |
| pH 7.2 vs pH 6.6 | 134771.1012 | 170153.2098 | 1.12014 | 0.99144 | 0.05 | n.s. | -445669.8465 | 715212.0489 |
| pH 7.2 vs pH 6.8 | 170031.6784 | 170153.2098 | 1.4132 | 0.96864 | 0.05 | n.s. | -410409.2694 | 750472.6261 |
| pH 7.2 vs pH 7.0 | 85739.83536 | 159163.7536 | 0.76182 | 0.99921 | 0.05 | n.s. | -457212.9545 | 628692.6253 |
| pH 7.4 vs pH 6.6 | 182549.8509 | 170153.2098 | 1.51725 | 0.95469 | 0.05 | n.s. | -397891.0968 | 762990.7986 |
| pH 7.4 vs pH 6.8 | 217810.4281 | 170153.2098 | 1.81031 | 0.89482 | 0.05 | n.s. | -362630.5196 | 798251.3758 |
| pH 7.4 vs pH 7.0 | 133518.5851 | 159163.7536 | 1.18635 | 0.98806 | 0.05 | n.s. | -409434.2048 | 676471.375 |
| pH 7.4 vs pH 7.2 | 47778.74975 | 170153.2098 | 0.39711 | 0.99999 | 0.05 | n.s. | -532662.198 | 628219.6975 |
| pH 7.6 vs pH 6.6 | 97086.77232 | 170153.2098 | 0.80693 | 0.99886 | 0.05 | n.s. | -483354.1754 | 677527.72 |
| pH 7.6 vs pH 6.8 | 132347.3495 | 170153.2098 | 1.09999 | 0.99231 | 0.05 | n.s. | -448093.5982 | 712788.2972 |
| pH 7.6 vs pH 7.0 | 48055.5065 | 159163.7536 | 0.42699 | 0.99998 | 0.05 | n.s. | -494897.2834 | 591008.2964 |
| pH 7.6 vs pH 7.2 | -37684.32886 | 170153.2098 | 0.31321 | 1 | 0.05 | n.s. | -618125.2766 | 542756.6189 |
| pH 7.6 vs pH 7.4 | -85463.07861 | 170153.2098 | 0.71032 | 0.9995 | 0.05 | n.s. | -665904.0263 | 494977.8691 |
| pH 7.8 vs pH 6.6 | -108448.3259 | 170153.2098 | 0.90136 | 0.99771 | 0.05 | n.s. | -688889.2737 | 471992.6218 |
| pH 7.8 vs pH 6.8 | -73187.74874 | 170153.2098 | 0.60829 | 0.99982 | 0.05 | n.s. | -653628.6965 | 507253.199 |
| pH 7.8 vs pH 7.0 | -157479.5918 | 159163.7536 | 1.39925 | 0.97025 | 0.05 | n.s. | -700432.3817 | 385473.1982 |
| pH 7.8 vs pH 7.2 | -243219.4271 | 170153.2098 | 2.0215 | 0.83244 | 0.05 | n.s. | -823660.3748 | 337221.5206 |
| pH 7.8 vs pH 7.4 | -290998.1769 | 170153.2098 | 2.41861 | 0.6814 | 0.05 | n.s. | -871439.1246 | 289442.7709 |
| pH 7.8 vs pH 7.6 | -205535.0983 | 170153.2098 | 1.70829 | 0.91924 | 0.05 | n.s. | -785976.046 | 374905.8495 |
| pH 8.0 vs pH 6.6 | -554992.2464 | 159163.7536 | 4.93126 | 0.04308 | 0.05 | significant | -1097950 | -12039.4565 |
| pH 8.0 vs pH 6.8 | -519731.6692 | 159163.7536 | 4.61796 | 0.06638 | 0.05 | n.s. | -1062680 | 23221.12069 |
| pH 8.0 vs pH 7.0 | -604023.5122 | 147357.0022 | 5.79693 | 0.01241 | 0.05 | significant | -1106700 | -101346.9061 |
| pH 8.0 vs pH 7.2 | -689763.3476 | 159163.7536 | 6.12874 | 0.00762 | 0.05 | significant | -1232720 | -146810.5577 |
| pH 8.0 vs pH 7.4 | -737542.0973 | 159163.7536 | 6.55326 | 0.00408 | 0.05 | significant | -1280490 | -194589.3074 |
| pH 8.0 vs pH 7.6 | -652079.0187 | 159163.7536 | 5.7939 | 0.01247 | 0.05 | significant | -1195030 | -109126.2288 |
| pH 8.0 vs pH 7.8 | -446543.9205 | 159163.7536 | 3.96767 | 0.15445 | 0.05 | n.s. | -989496.7104 | 96408.86944 |

#### Table S2

**Analysis of variance with Turkey's post-hoc test on the data from low-speed co-sedimentation of F-actin in the presence of PLS2 with varying pH.** Fraction of F-actin pelleted using a low-speed co-sedimentation assay in each pH condition compared to each other pH condition; n.s. is non-significant.
Related to Figure 1D.

| pH condition | Δ mean | SE of mean | q Value | Probability | Alpha | Significance | Lower control limit | Upper control limit |
|---|---|---|---|---|---|---|---|---|
| pH 6.8 vs pH 6.6 | 0.13089 | 0.07612 | 2.43173 | 0.67619 | 0.05 | n.s. | -0.13266 | 0.39444 |
| pH 7.0 vs pH 6.6 | 0.21019 | 0.07612 | 3.90494 | 0.1736 | 0.05 | n.s. | -0.05336 | 0.47374 |
| pH 7.0 vs pH 6.8 | 0.0793 | 0.07612 | 1.47321 | 0.96021 | 0.05 | n.s. | -0.18425 | 0.34285 |
| pH 7.2 vs pH 6.6 | 0.19703 | 0.07612 | 3.66048 | 0.2291 | 0.05 | n.s. | -0.06652 | 0.46058 |
| pH 7.2 vs pH 6.8 | 0.06614 | 0.07612 | 1.22874 | 0.98501 | 0.05 | n.s. | -0.19741 | 0.32969 |
| pH 7.2 vs pH 7.0 | -0.01316 | 0.07612 | 0.24447 | 1 | 0.05 | n.s. | -0.27671 | 0.25039 |
| pH 7.4 vs pH 6.6 | 0.20498 | 0.07612 | 3.80801 | 0.19414 | 0.05 | n.s. | -0.05857 | 0.46853 |
| pH 7.4 vs pH 6.8 | 0.07408 | 0.07612 | 1.37628 | 0.97211 | 0.05 | n.s. | -0.18947 | 0.33763 |
| pH 7.4 vs pH 7.0 | -0.00522 | 0.07612 | 0.09693 | 1 | 0.05 | n.s. | -0.26877 | 0.25833 |
| pH 7.4 vs pH 7.2 | 0.00794 | 0.07612 | 0.14754 | 1 | 0.05 | n.s. | -0.25561 | 0.27149 |
| pH 7.6 vs pH 6.6 | 0.20107 | 0.07612 | 3.73538 | 0.21078 | 0.05 | n.s. | -0.06248 | 0.46462 |
| pH 7.6 vs pH 6.8 | 0.07017 | 0.07612 | 1.30365 | 0.97919 | 0.05 | n.s. | -0.19338 | 0.33372 |
| pH 7.6 vs pH 7.0 | -0.00913 | 0.07612 | 0.16956 | 1 | 0.05 | n.s. | -0.27268 | 0.25442 |
| pH 7.6 vs pH 7.2 | 0.00403 | 0.07612 | 0.07491 | 1 | 0.05 | n.s. | -0.25952 | 0.26758 |
| pH 7.6 vs pH 7.4 | -0.00391 | 0.07612 | 0.07263 | 1 | 0.05 | n.s. | -0.26746 | 0.25964 |
| pH 7.8 vs pH 6.6 | 0.09868 | 0.07612 | 1.83323 | 0.88754 | 0.05 | n.s. | -0.16487 | 0.36223 |
| pH 7.8 vs pH 6.8 | -0.03222 | 0.07612 | 0.5985 | 0.99983 | 0.05 | n.s. | -0.29577 | 0.23133 |
| pH 7.8 vs pH 7.0 | -0.11151 | 0.07612 | 2.07171 | 0.81437 | 0.05 | n.s. | -0.37506 | 0.15204 |
| pH 7.8 vs pH 7.2 | -0.09836 | 0.07612 | 1.82725 | 0.88913 | 0.05 | n.s. | -0.36191 | 0.16519 |
| pH 7.8 vs pH 7.4 | -0.1063 | 0.07612 | 1.97478 | 0.84632 | 0.05 | n.s. | -0.36985 | 0.15725 |
| pH 7.8 vs pH 7.6 | -0.10239 | 0.07612 | 1.90215 | 0.86832 | 0.05 | n.s. | -0.36594 | 0.16116 |
| pH 8.0 vs pH 6.6 | -0.05992 | 0.07612 | 1.11311 | 0.9915 | 0.05 | n.s. | -0.32347 | 0.20363 |
| pH 8.0 vs pH 6.8 | -0.19081 | 0.07612 | 3.54484 | 0.25975 | 0.05 | n.s. | -0.45436 | 0.07274 |
| pH 8.0 vs pH 7.0 | -0.27011 | 0.07612 | 5.01805 | 0.04252 | 0.05 | significant | -0.53366 | -0.00656 |
| pH 8.0 vs pH 7.2 | -0.25695 | 0.07612 | 4.77359 | 0.05877 | 0.05 | n.s. | -0.5205 | 0.0066 |
| pH 8.0 vs pH 7.4 | -0.26489 | 0.07612 | 4.92112 | 0.04838 | 0.05 | significant | -0.52844 | -0.00134 |
| pH 8.0 vs pH 7.6 | -0.26098 | 0.07612 | 4.84849 | 0.05326 | 0.05 | n.s. | -0.52453 | 0.00257 |
| pH 8.0 vs pH 7.8 | -0.15859 | 0.07612 | 2.94634 | 0.46326 | 0.05 | n.s. | -0.42214 | 0.10496 |

### Table S3

**Kinetic and equilibrium parameters for association and dissociation of FM-ABD2 to and from RD-ABD1 at varying buffer pH.** $k_{off}$ ($n \geq 4$) and $K_d$ ($n = 3$) measured in equilibrium assays are reported as mean ± SD. Error for $k_{on}$ represents curve fit error. $K_d$ derived from kinetics was calculated using the $k_{off}$ and $k_{on}$ for each specified pH.
Related to Figure 3.

| pH | $k_{on}$ | $k_{off}$ | $K_d$ derived from kinetics (nM) | $K_d$ from equilibrium (nM) |
|---|---|---|---|---|
| 6.6 | 1.94 ± 0.02 | 0.010 ± 0.003 | 5.1 | – |
| 7.0 | 1.60 ± 0.02 | 0.009 ± 0.002 | 5.7 | 4.5 ± 0.3 |
| 7.4 | 1.24 ± 0.02 | 0.013 ± 0.002 | 11 | – |
| 7.8 | 1.07 ± 0.02 | 0.008 ± 0.001 | 7.7 | – |
| 8.0 | 1.010 ± 0.008 | 0.008 ± 0.001 | 7.6 | 8.7 ± 0.8 |

#### Table S4

**Analysis of variance with Turkey's post-hoc test on the data from high-speed co-sedimentation experiments.**
n.s. is non-significant.
Related to Figure 4D.

| pH condition | Δ mean | SE of mean | q Value | Probability | Alpha | Significance | Lower control limit | Upper control limit |
|---|---|---|---|---|---|---|---|---|
| ABD1 pH 7 vs. ABD1 pH 8 | 0.2106 | 0.03514 | 8.477 | 0.0007 | 0.05 | significant | 0.09260 | 0.3286 |
| ABD1 pH 7 vs. ABD1 H207K pH 7 | 0.01124 | 0.03514 | 0.4525 | 0.9994 | 0.05 | ns | -0.1068 | 0.1293 |
| ABD1 pH 7 vs. ABD1 H207K pH 8 | 0.3031 | 0.03514 | 12.20 | <0.0001 | 0.05 | significant | 0.1851 | 0.4211 |
| ABD1 pH 7 vs. ABD1 H207Y pH 7 | 0.1379 | 0.03514 | 5.552 | 0.0192 | 0.05 | significant | 0.01992 | 0.2560 |
| ABD1 pH 7 vs. ABD1 H207Y pH 8 | 0.2935 | 0.03514 | 11.81 | <0.0001 | 0.05 | significant | 0.1754 | 0.4115 |
| ABD1 pH 8 vs. ABD1 H207K pH 7 | -0.1994 | 0.03514 | 8.025 | 0.0011 | 0.05 | significant | -0.3174 | -0.08136 |
| ABD1 pH 8 vs. ABD1 H207K pH 8 | 0.09247 | 0.03514 | 3.722 | 0.1625 | 0.05 | ns | -0.02555 | 0.2105 |
| ABD1 pH 8 vs. ABD1 H207Y pH 7 | -0.07268 | 0.03514 | 2.925 | 0.3627 | 0.05 | ns | -0.1907 | 0.04534 |
| ABD1 pH 8 vs. ABD1 H207Y pH 8 | 0.08285 | 0.03514 | 3.335 | 0.2444 | 0.05 | ns | -0.03517 | 0.2009 |
| ABD1 H207K pH 7 vs. ABD1 H207K pH 8 | 0.2918 | 0.03514 | 11.75 | <0.0001 | 0.05 | significant | 0.1738 | 0.4099 |
| ABD1 H207K pH 7 vs. ABD1 H207Y pH 7 | 0.1267 | 0.03514 | 5.099 | 0.0330 | 0.05 | significant | 0.008675 | 0.2447 |
| ABD1 H207K pH 7 vs. ABD1 H207Y pH 8 | 0.2822 | 0.03514 | 11.36 | <0.0001 | 0.05 | significant | 0.1642 | 0.4002 |
| ABD1 H207K pH 8 vs. ABD1 H207Y pH 7 | -0.1652 | 0.03514 | 6.647 | 0.0053 | 0.05 | significant | -0.2832 | -0.04713 |
| ABD1 H207K pH 8 vs. ABD1 H207Y pH 8 | -0.009622 | 0.03514 | 0.3873 | 0.9997 | 0.05 | ns | -0.1276 | 0.1084 |
| ABD1 H207Y pH 7 vs. ABD1 H207Y pH 8 | 0.1555 | 0.03514 | 6.260 | 0.0083 | 0.05 | significant | 0.03751 | 0.2735 |

#### Table S5

**Kinetic parameters for association and dissociation of FM-ABD2 to and from RD-ABD1 variants at pH 7.0 and 8.0.** $k_{off}$ values reported as mean ± SD (n ≥ 4). Error for $k_{on}$ represents curve fit error. $K_d$ derived from kinetics was calculated using the $k_{off}$ and $k_{on}$ for the specified pH and construct. Since the dissociation rate of RD-ABD1-H207K at pH 8.0 followed a double exponential fit with 62% and 38% amplitudes (A), two corresponding $k_{off}$ values are reported.
Related to Figure 4E,F.

| | | $k_{on}$ | $k_{off}$ | $K_d$ derived from kinetics (nM) |
|---|---|---|---|---|
| RD-ABD1 | pH 7.0 | 1.60 ± 0.02 | 0.009 ± 0.002 | 5.7 |
| | pH 8.0 | 1.01 ± 0.01 | 0.008 ± 0.001 | 7.6 |
| RD-ABD1-H207K | pH 7.0 | 1.59 ± 0.02 | 0.039 ± 0.001 | 24 |
| | pH 8.0 | 1.12 ± 0.02 | 0.097 ± 0.004 (0.62 A) | 87 |
| | | | 0.016 ± 0.002 (0.38 A) | 14 |
| RD-ABD1-H207Y | pH 7.0 | 1.87 ± 0.02 | 0.0021 ± 0.0001 | 1.1 |
| | pH 8.0 | 1.21 ± 0.01 | 0.0026 ± 0.0004 | 2.2 |

### Supplementary Video Captions

Videos S1-S4 are related to Figure 5.

**Video S1.** Time-lapse imaging of cells transiently co-transfected with mCherry-β-actin and mEmerald-PLS2-WT sequentially clamped at the indicated pH values.

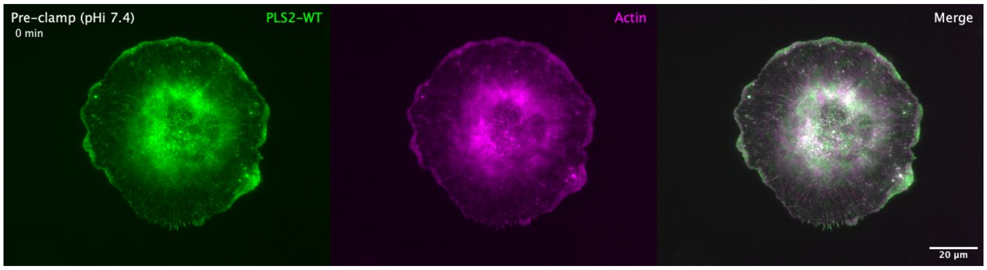


**Video S2.** Time-lapse imaging of cells transiently co-transfected with mCherry-β-actin and mEmerald-PLS2-S5D sequentially clamped at the indicated pH values.

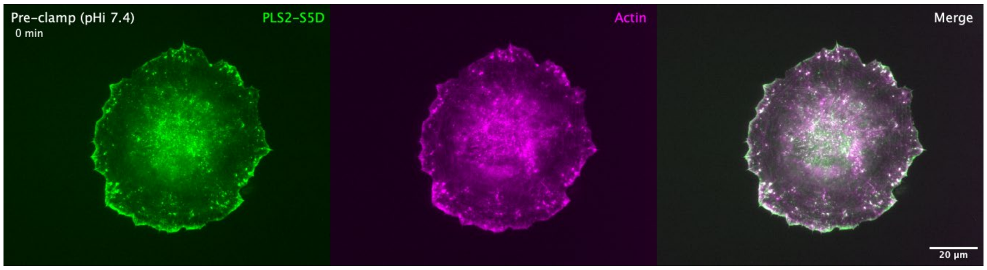


**Video S3.** Time-lapse imaging of cells transiently co-transfected with mCherry-β-actin and mEmerald-PLS2-S5D/H207K sequentially clamped at the indicated pH values.

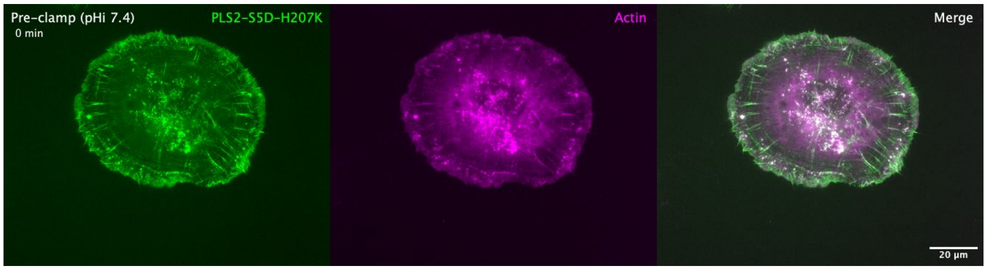


**Video S4.** Time-lapse imaging of cells transiently co-transfected with mCherry-β-actin and mEmerald-PLS2-S5D/H207Y sequentially clamped at the indicated pH values.

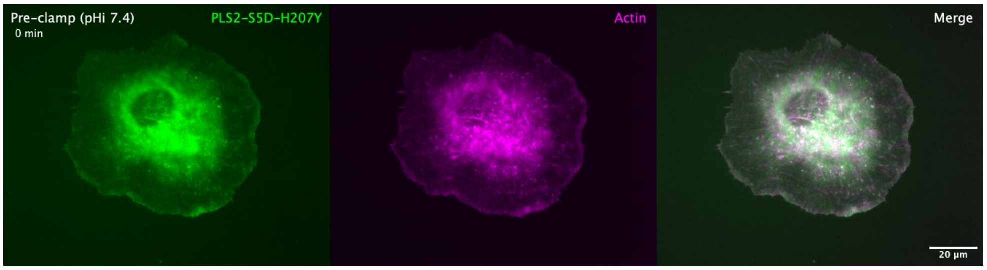
